# TET2 loss impairs MPLA-induced innate immune memory during infection and disrupts hematopoiesis via RIPK1 in mice

**DOI:** 10.1101/2025.09.08.674943

**Authors:** Alyssa N. L. Jarabek, Xenia Davis, Mary Oliver, Yongchao Wang, Heidi Chen, Julia K. Bohannon, Sandra Zinkel

## Abstract

Ten-eleven translocation protein 2 (TET2) is commonly mutated in hematologic disorders of bone marrow failure such as myelodysplastic syndrome (MDS), and loss of TET2 is associated with poor prognosis. TET2 loss is also associated with augmented inflammation, as well as impaired innate immune responses. However, many individuals harboring mutations do not develop hematologic disorders, indicating that additional factors, such as inflammation, cooperate with TET2 loss to promote disease progression. Innate immune memory is a phenomenon in which pre-treatment with microbial ligands, including monophosphoryl lipid A (MPLA), improves innate immune function while minimizing inflammation during subsequent infection. Given the previously reported innate immune impairments attributed to TET2 loss, we tested whether MPLA could induce an innate immune response in TET2-deficient mice infected with *P. aeruginosa*. Moreover, due to the inflammatory role of TET2 loss, we investigated whether MPLA with infection would promote disease-related phenotypes. We found that TET2-deficient mice display impaired infection response to *P. aeruginosa* which is partially improved with MPLA pretreatment. Assessments of pathogenic clearance functions further showed decreased capacity in TET2-deficient innate immune cells. Moreover, TET2-defcient cells also exhibit impaired differentiation. MPLA pretreatment in infected mice promotes myeloid-biased hematopoiesis at the expense of erythroid- and megakaryocyte-biased hematopoiesis, resembling MDS-like disease progression. Intriguingly, inhibition of receptor-interacting serine/threonine-protein kinase 1 (RIPK1) dampens the effects of TET2 loss on hematopoietic perturbation. Collectively, we find that TET2 loss impairs innate immune memory and infection responses, and MPLA with infection promotes hematopoietic skewing through RIPK1.

**KEY POINTS:**

- TET2 loss increases susceptibility to infection, but immune response can be improved by MPLA-induced innate immune memory.
- MPLA pretreatment and infection promotes aberrant hematopoiesis and myeloid bias in TET2 deficiency that is improved by RIPK1 inhibition.

## INTRODUCTION

Hematopoiesis is a dynamic process, designed to appropriately respond to environmental stresses to maintain proper blood cell production. The entire system originates from hematopoietic stem cells (HSCs) that sense and respond to stress by altering and skewing blood cell production to craft a stress-appropriate response. During aging, HSCs acquire mutations, and those that confer a survival or proliferation advantage are propagated through hematopoiesis, a process known as clonal hematopoiesis (CH).^1,2^ The presence of CH imparts increased risk of developing hematopoietic diseases such as myelodysplastic syndrome (MDS), acute myelogenous leukemia (AML), chronic myelomonocytic leukemia (CMML), cardiovascular disease, and a variety of other diseases. However, most individuals with CH do not develop disease, suggesting that additional genetic and/or environmental factors contribute to disease development.^3,4^

Epigenetic regulators such as <u>T</u>en-<u>e</u>leven translocation protein <u>2</u> (TET2) and <u>DN</u>A <u>m</u>ethyltransferase <u>3a</u> (DNMT3a), are the most commonly mutated genes in CH. DNMT3a and TET2 are initiating mutations that causally associate with hematopoietic and cardiovascular diseases. TET2 is a cytosine demethylase that catalyzes the conversion of 5-methylcytosine to 5-hydroxymethylcytosine and commonly exhibits loss-of-function mutation in CH and MDS.^5–8^ TET2 is also involved in innate immune responses. *TET2*^-/-^ in human neutrophils impairs phagocytosis and NETosis, and TET2-deficient mice exhibit decreased survival and heightened pathology during *Streptococcal pneumoniae* infection.^9,10^ Moreover, TET2 tightly regulates inflammatory responses.^11–15^ TET2-deficient mice have heightened and impaired resolution of inflammation following lipopolysaccharide (LPS).^12^ Additionally, LPS-induced inflammation promotes *TET2-*deficient HSC expansion, which is partially mitigated by inflammatory pathway inhibition.^12–15^ Thus, inflammatory challenge confers a survival and proliferation advantage to TET2-deficient HSCs in a feed-forward manner.

Infection is one of the most common environmental inflammatory challenges encountered during an individual’s lifetime. Bacterial infections activate the innate immune system and induce epigenetic and metabolic rewiring of hematopoietic stem and progenitors and macrophages that potentiate innate immune responses to subsequent infections in a process known as innate immune memory.^16–23^ HSCs are reservoirs of innate immune memory, and, along with macrophages, direct enhanced responses and cross-protection against subsequent infections.^17,24–26^ Enhanced responses can include augmented bacterial clearance, phagocytosis, reactive oxygen species (ROS) production, leukocyte recruitment, survival protection, and altered cytokine production.^16,18,19,23,27–29^ Depending on the inciting stimulus, innate immune memory elicits protection on the order of weeks to months, distinguishing this form of immunological memory from adaptive protection.^24,29^ Collectively, these functions control and clear bacteria to prevent severe infection and expedite recovery.

In addition to infection, innate immune memory is elicited by a variety of substances derived from infectious agents or pathogen-associated molecular patterns (PAMPS), such as fungal cell walls (β-Glucan), or bacterial cell walls (peptidoglycan, Monophosphoryl lipid A, or MPLA). MPLA is a toll-like receptor 4 (TLR4) agonist and non-toxic derivative of LPS that induces innate immune memory and dampens pro-inflammatory cytokine responses following secondary stimulus, conferring potential to prevent pathological consequences of inflammatory cytokines that contribute to morbidity and mortality.^16,19,26,28,29^ On the other hand, data from mouse models demonstrates HSC function is compromised following bacterial infection, an effect mediated by TLR4-direct pathogen sensing by HSCs.^30^

Given the detrimental effect of inflammation on HSC function, strategies to interrupt this signaling warrant attention. RIPK1 (<u>R</u>eceptor-interacting serine/threonine-<u>p</u>rotein <u>k</u>inase <u>1</u>) is pharmacologically targetable and a mediator of inflammatory signaling downstream of death receptors and TLR4.^31^ RIPK1 also mediates inflammatory cell death (necroptosis) in hematopoietic stem and progenitors.^32,33^ RIPK1 and necroptosis signaling is increased in human MDS patient bone marrow (BM) samples, most prominent in the erythroid islands.^34^ During TLR4 signaling, RIPK1 binds to TIR domain-containing adaptor protein inducing IFN-β (TRIF) to activate interferon responses, NFκB, and pro-inflammatory gene expression.^31^ RIPK1 is therefore situated at the nexus of critical signaling pathways directing HSC responses to pathogens and cytokines.

TET2 loss impairs innate immune responses, which hinders bacterial clearance and prolongs infection and inflammation. TET2 loss also impairs resolution of inflammation and inflammatory signaling through TLR4 that promotes HSC expansion and myeloid-biased hematopoiesis.^12^ Bacterial infection of individuals harboring *Tet2-*inactivation could therefore promote clonal expansion and disease progression.^35^ Innate immune function augmentation through inducing innate immune memory has potential to improve response to infection, but may promote HSC dysfunction through TLR4-TRIF signaling. Thus, we tested whether MPLA induces innate immune memory in *VavTet2-*deficient (*vavTet2^F/F^*) mice and improves innate immune response to *Pseudomonas aeruginosa* infection. We further tested whether MPLA treatment prior to infection exacerbates hematopoietic stem and progenitor abnormalities in *VavTet2^F/F^* mice, and whether hematopoietic dysfunction is dampened by genetic RIPK1 kinase inhibition (*VavTet2^F/F^Ripk1^D^*^138^*^N/+^*).^36^

We found that *VavTet2^F/F^* mice display exacerbated response to *P. aeruginosa* infection, and MPLA- pretreated *VavTet2^F/F^*mice have suboptimal protection from severe infection and impaired innate immune function. MPLA administration prior to *P. aeruginosa* promotes myeloid-biased hematopoiesis and expands HSCs in *VavTet2^F/F^* mice. However, RIPK1 kinase inactivation dampens the effects of MPLA pretreatment with *P. aeruginosa* on hematopoietic dysfunction in *VavTet2^F/F^* mice. These results show that *VavTet2^F/F^* mice develop innate immune memory, although suboptimal. Additionally, MPLA administration prior to *P. aeruginosa* infection promotes myeloid-biased hematopoiesis and HSC expansion that is dampened by RIPK1 kinase inactivation. These results reveal the effect of *Tet2* loss on innate immune memory and response to infection. This study further elucidates the impact of infection and innate immune memory on hematopoiesis in *VavTet2^F/F^* mice and demonstrates a potential role for RIPK1 inhibition in dampening its detrimental effects on hematopoiesis.

## MATERIALS AND METHODS

### Mice

*vavCreTet2^fl/fl^* mice (hereafter *VavTet2^F/F^*) and *vavCreTet2^fl/fl^Ripk1^D^*^138^*^N/+^* mice (hereafter *VavTet2^F/F^Ripk1^D/+^*) were generated from *Tet2^fl/fl^* mice and *Ripk1^D^*^138^*^N/D^*^138^*^N^* mice, obtained from Omar Abdel-Wahab and Michelle Kelliher, respectively.^36,37^ Both male and female mice, aged 6-34 weeks were used, age- and sex-matched across genotypes and treatment groups. The Vanderbilt University Institutional Animal Care and Use Committee approved all experiments (#V1800188-02 and M1800068-02). Complete description in Supplemental Methods.

### Cell Culture

BM was harvested as previously described and cultured in complete media with 10% L929-conitioned media for 7 days.^38^ MPLA and LPS were used at concentrations indicated in figure legends. Complete description in Supplemental Methods.

### Flow Cytometry of Infected Murine Bone Marrow and Peritoneal Cells

Red cell lysis was performed on bone marrow (BM), and both BM and peritoneal fluid (PF) were filtered and incubated with fluorochrome-conjugated antibodies (Supplemental Table 3). Cells were pelleted and incubated with 2% paraformaldehyde prior to assessment via BD Cytek® Aurora (Cytek® Biosciences, Fremont, California). Complete description in Supplemental Methods.

### Statistical analysis

Statistics were performed using GraphPad Prism software, version 10.4.1 (La Jolla, California). Outliers were identified and excluded using the ROUT test. Data were analyzed using either one- or two-way ANOVA with post-hoc Tukey’s test, as indicated in figure legends. Data are mean ± standard error of the mean (SEM), unless otherwise indicated. Flow cytometric data was analyzed using Flowjo software (Ashland, OR) and OMIQ software platform using cyCombine for batch normalization across multiple runs (Dotmatics, Boston, MA).

## RESULTS

### *Tet2* loss exacerbates LPS-induced inflammatory signaling through RIPK1

*Tet2* loss upregulates pro-inflammatory cytokine production during LPS stimulation *in vitro* and *in vivo*.^11,12^ Additionally, RIPK1 is essential for inflammatory signaling.^39,40^ Therefore, we investigated how hematopoietic-specific loss of *Tet2* during LPS *in vivo* affects inflammation and the role of RIPK1 in this phenomenon. We treated WT, *VavTet2^F/F^*, and *VavTet2^F/F^Ripk1^D/+^* mice with LPS intraperitoneally and isolated RNA from whole BM 8 hours later. NanoString analysis of myeloid innate immunity-related transcripts revealed that LPS-treated *VavTet2^F/F^* mice but not WT or *VavTet2^F/F^Ripk1^D/+^* mice upregulate inflammatory and immune-related genes (Figure 1A). Principal component analysis (PCA) also revealed that the NanoString profile of LPS-treated *VavTet2^F/F^* mice segregates independently of other groups (Figure 1B). Gene differential expression analysis indicates upregulation of *Tnfa*, *Il1b*, and *Il6* following LPS treatment in *VavTet2^F/F^* mice compared to WT or *VavTet2^F/F^Ripk1^D/+^* (Figure 1C). RT-qPCR on BM from PBS- and LPS-treated mice further validates expression of inflammatory genes (Figure 1D-E, Supplemental Figure 1A). Collectively, these data demonstrate that *Tet2* loss exacerbates inflammatory response to LPS-induced stress *in vivo* in a RIPK1- dependent manner.

**Figure 1:**
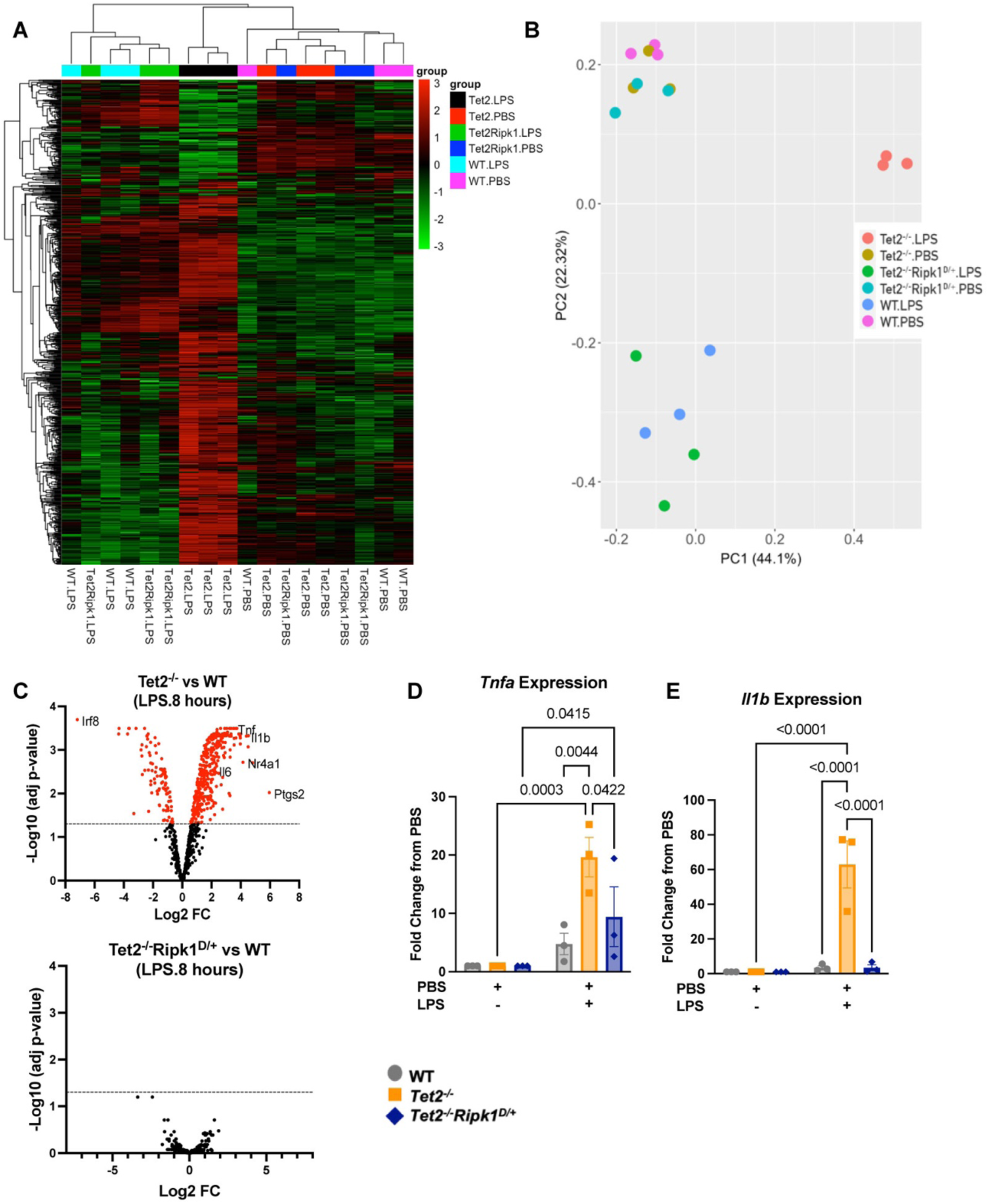
*Tet2* loss exacerbates LPS-induced inflammatory signaling through RIPK1. WT, *Tet2^-/-^*, and *Tet2Ripk1^D/+^* mice were treated with PBS or LPS (1.5 mg/kg) for 8 hours, followed by bone marrow collection, RNA isolation and RNA profiling of immunity-related genes by NanoString analysis (n=3). **(A)** Spearman correlation heatmap generated and clustered based on gene expression in log2 scale. **(B)** PCA plot Principal component analysis (PCA) on log2 transformed gene expression. **(C)** Gene differential expression analysis in the bone marrow between LPS-treated WT and *Tet2^-/-^* mice (upper) or LPS-treated WT and *Tet2Ripk1^D/+^* mice (lower). **(D-E)** Validation of the expression of inflammatory genes (*Tnfa* and *Il1b*) by qRT-PCR. Data are shown as mean ± SEM two-way ANOVA with post-hoc Tukey’s test, n=3 mice per group. Each point represents an individual mouse.

#### *Tet2*-deficient mice display impaired response to *P. aeruginosa* infection

To determine if *Tet2* loss and RIPK1 inactivation impairs bacterial infection response and innate immune protection, we intraperitoneally (IP) injected WT, *VavTet2^F/F^*, and *VavTet2^F/F^Ripk1^D/+^* mice with vehicle (lactated Ringers, LR) or MPLA, and 3 days later, infected with *P. aeruginosa* or LR *via* IP injection (Figure 2A). At 6- hours post-infection (pi), we assessed immune responses. LR-treated *VavTet2^F/F^* and *VavTet2^F/F^Ripk1^D/+^*mice display decreased body temperature compared to WT LR-treated mice, consistent with impaired infection response (Figure 2B). Notably, MPLA pre-treatment in all genotypes significantly recovers body temperatures to baseline averages (Figure 2B). All genotypes decrease bacterial burden in PF with MPLA pre-treatment (Figure 2C). When bacterial burden is examined in MPLA-treated groups separately, CFU/mL in *VavTet2^F/F^* mice is significantly higher than WT (Figure 2D). These data demonstrate that *VavTet2^F/F^* and *VavTet2^F/F^Ripk1^D/+^* mice can develop MPLA-induced innate immune memory. However, *VavTet2^F/F^* mice impair bacterial clearance relative to WT mice.

**Figure 2:**
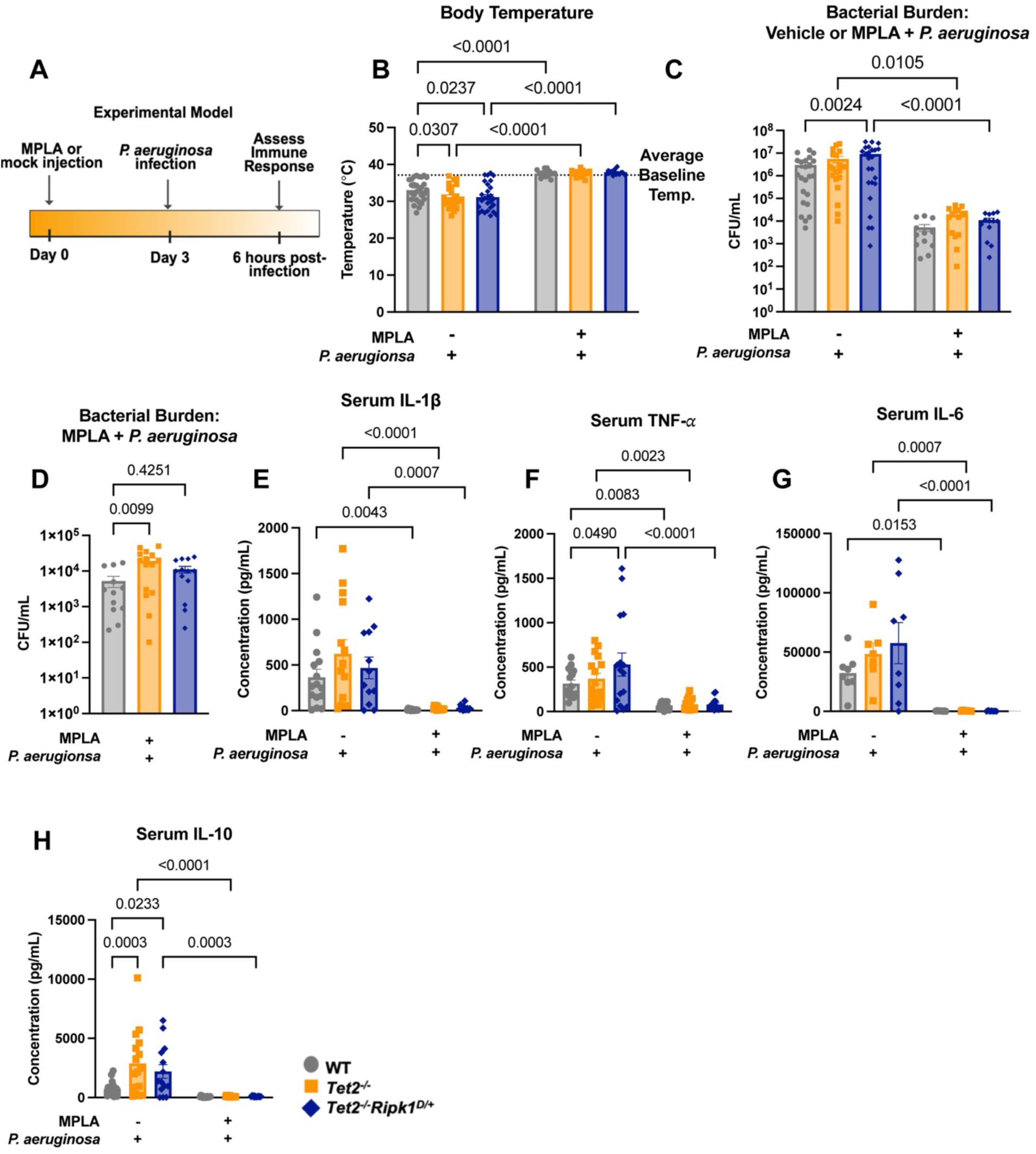
*Tet2*-deficient mice display impaired response to *P. aeruginosa* infection. **(A)** Schematic of experimental design. Mice were intraperitoneally injected with MPLA (20 µg) or LR at day 0. 3 days post-injection, mice were infected with *P. aeruginosa* or LR for 6 hours prior to immune response assessment. **(B)** Rectal body temperature was assessed at 6 hr. post-infection, and **(C-D)** peritoneal fluid was harvested for bacterial burden assessment. **(E-H)** Plasma was collected from mice, and Bio-Plex assay was performed to determine concentration of circulating serum cytokines. Data are shown as mean ± SEM, one- or two-way ANOVA with post-hoc Tukey’s test. n = 15-28 mice per group for body temperature and bacterial burden, n = 15-16 mice per group for serum cytokines. Each point represents an individual mouse.

### MPLA administered prior to infection dampens proinflammatory cytokines

To elucidate the systemic inflammatory response to *P. aeruginosa* infection, we examined serum cytokines in infected and MPLA-pretreated mice. MPLA pretreatment reduces proinflammatory cytokines in all genotypes during infection, though LR-pretreated *VavTet2^F/F^* mice display a trend towards increased proinflammatory cytokines relative to WT (Figure 2E-G). Additionally, *VavTet2^F/F^* and *VavTet2^F/F^Ripk1^D/+^* LR-pretreated mice increase IL-10 compared to WT, consistent with a compensatory effect of augmented proinflammatory cytokines (Figure 2H). When compared to uninfected mice given LR or MPLA at 3 days post-administration, the magnitude of cytokine production is low compared to infected counterparts (Supplemental Figure 1B-G). Intriguingly, *VavTet2^F/F^* mice reduce granulocyte colony stimulating factor (G-CSF) in MPLA-treated groups compared to untreated, yet all genotypes show reductions with MPLA and infection, likely due to reduced bacterial load (Supplemental Figure 1H-I). Granulocyte-macrophage colony stimulating factor (GM-CSF) decreases in *VavTet2^F/F^*and *VavTet2^F/F^Ripk1^D/+^* mice pretreated with MPLA compared to WT, and in MPLA-pretreated and infected mice compared to infected only groups (Supplemental Figure 1J-K). Altogether, these data demonstrate that *VavTet2^F/F^* mice can respond to MPLA pre-treatment to significantly reduce circulating pro-inflammatory cytokines during infection.

To examine the cell-specific effect of macrophages on the inflammatory effects of *Tet2* loss in MPLA-induced innate immune memory, we stimulated BM-derived macrophages (BMDMs) with MPLA for 3 days and challenged with LPS prior to proinflammatory gene expression assessment. Both *VavTet2^F/F^* and *VavTet2^F/F^Ripk1^D/+^* BMDMs increase pro-inflammatory gene expression compared to WT with either LPS alone or with both MPLA and LPS (Supplemental Figure 1L-M). These results suggest that the BM niche plays an important role in regulating the effect of RIPK1 dampening of pro-inflammatory effects of *Tet2* loss.

### *VavTet2^F/F^* mice recruit comparable populations of innate immune populations to infection site

To determine the mechanism of impaired bacterial clearance and innate immune protection in *Tet2* loss, we examined innate immune populations recruited to the infection site by flow cytometry analysis of PF. Mice of all genotypes increase peritoneal macrophages in MPLA-pretreated groups compared to LR-treated controls, consistent with effective innate immune memory responses (Figure 3A, Supplemental Figure 2A-B). However, *VavTet2^F/F^* mice but not WT or *VavTet2^F/F^Ripk1^D/+^* mice display impaired fold change macrophage frequency from LR-pretreated to MPLA-pretreated (Figure 3B). Ly6G^high^ neutrophils (mature populations) undergo a similar effect; mature neutrophils in WT, *VavTet2^F/F^*, and *VavTet2^F/F^Ripk1^D/+^* mice significantly increase with MPLA pre-treatment, yet fold change of Ly6G^high^ neutrophils from vehicle- to MPLA-pretreated is dampened in *VavTet2^F/F^* mice (Figure 3C-D, Supplemental Figure 2C-D). Less mature, Ly6G^mid^ neutrophils in WT and *VavTet2^F/F^Ripk1^D/+^*but not *VavTet2^F/F^* increase in frequency, which is similarly reflected in fold change frequency from vehicle-pretreated to MPLA-pretreated (Figure 3E-F, Supplemental Figure 2E-F). Thus, *VavTet2^F/F^* mice recruit innate cells to the infection site with MPLA-pretreatment, however there is diminished fold-change frequency following MPLA and decreased less-mature neutrophil populations.

**Figure 3:**
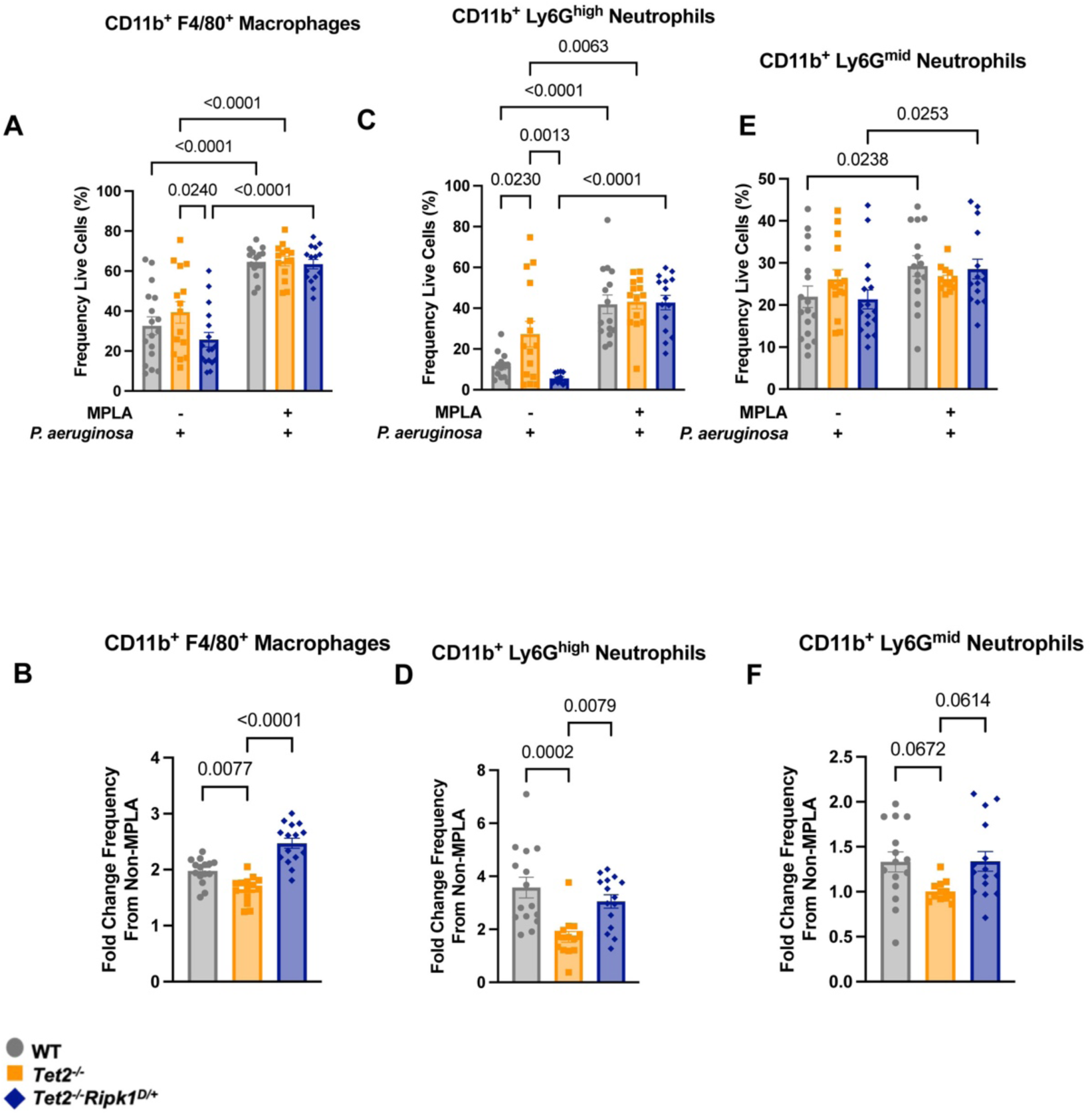
*VavTet2^F/F^* mice recruit comparable populations of innate immune populations to infection site. Peritoneal fluid was harvested from vehicle or MPLA-pretreated mice infected with *P. aeruginosa*. **(A)** Macrophages (CD11b^+^ F4/80^+^), as frequency of live cells were quantified, and **(B)** fold change frequency from non-MPLA-pretreated to MPLA pre-treated was quantified. **(C)** Mature neutrophils (CD11b^+^ Ly6G^high^), as frequency of live cells, and **(D)** fold change frequency non-MPLA- to MPLA-pretreated. **(E)** Less mature neutrophils (CD11b^+^ Ly6G^mid^) were quantified as frequency live cells, and **(F)** fold change frequency non-MPLA- to MPLA-pretreated. Data are shown as mean ± SEM, one- or two- way ANOVA with post-hoc Tukey’s test, n = 14-17 mice per group. Each point represents an individual mouse.

### *VavTet2^F/F^* innate immune cells are impaired in pathogenic clearance functions

As *VavTet2^F/F^* mice recruit comparable innate immune populations infection sites, we evaluated whether bacterial clearance deficits are due to pathogenic clearance functions. Total BM from *VavTet2^F/F^* mice in both LR- and MPLA-pretreated and infected groups increase cytosolic ROS compared to WT (Figure 4A). This trend is preserved in specific innate immune populations (Figure 4B-4C, Supplemental Figure 3A). Interestingly, *VavTet2^F/F^* and *VavTet2^F/F^Ripk1^D/+^* mice produce less ROS from LR-pretreatment to MPLA-pretreatment at the infection site in PF macrophages, neutrophils, and monocytes (Figure 4D-E, Supplemental Figure 3B-C). *In vivo* phagocytosis assay (Figure 4F) reveals that MPLA pre-treatment increases phagocytic total cell number in macrophages from all genotypes (Figure 4G). Interestingly, *VavTet2^F/F^* macrophages fail to augment phagocytic capacity per cell (geometric MFI, gMFI; Figure 4H). Moreover, *VavTet2^F/F^* and *VavTet2^F/F^Ripk1^D/+^* total neutrophil phagocytic count fail to augment with MPLA (Figure 4I, Supplemental Figure 3D). No notable changes occurred in monocytic phagocytosis capacity (Supplemental Figure 3E-F). Altogether, these results suggest that innate immune cell populations recruited to the infection site are impaired in pathogenic clearance functions in *VavTet2^F/F^* and *VavTet2^F/F^Ripk1^D/+^*mice, consistent with observed deficits in bacterial clearance.

**Figure 4:**
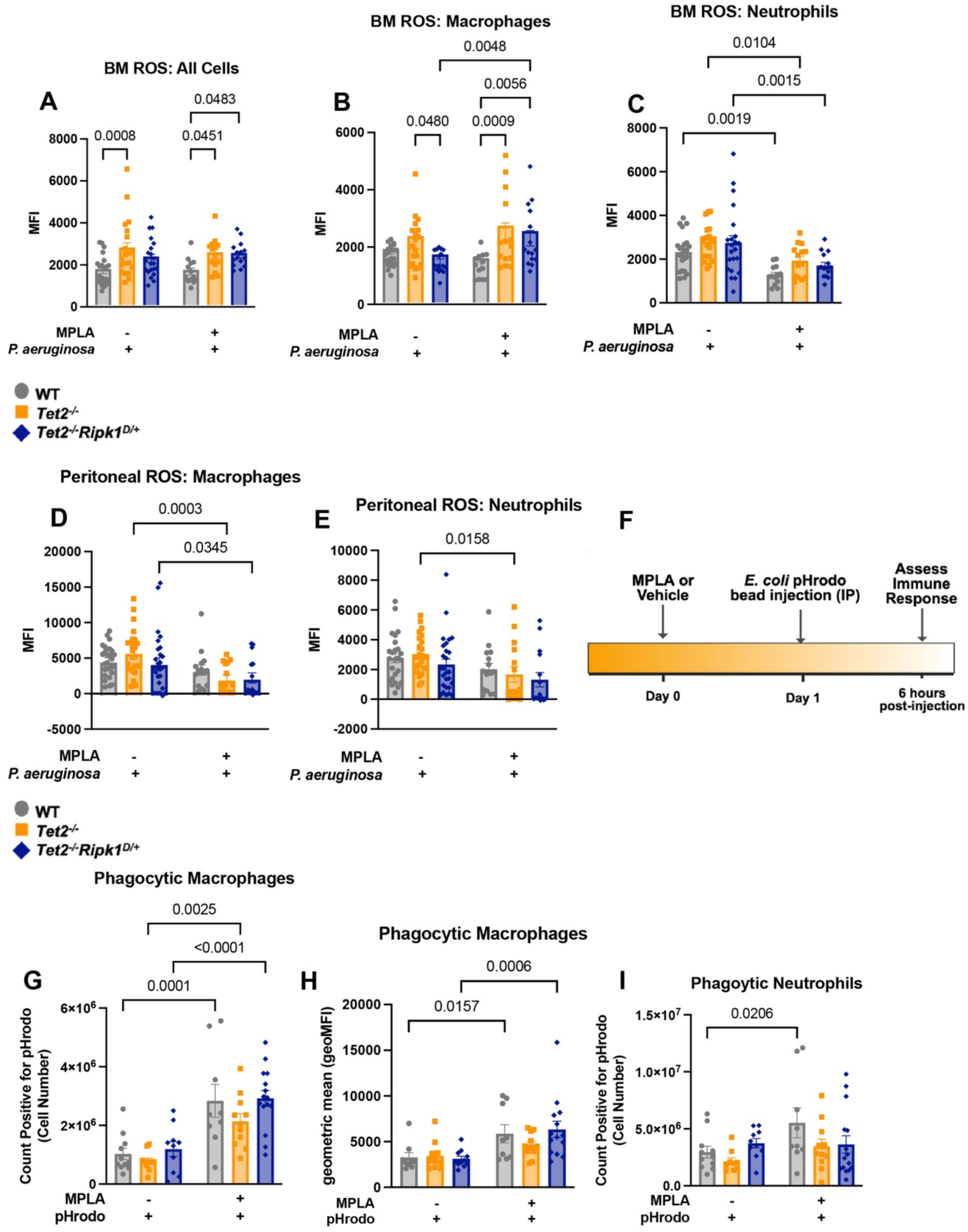
*VavTet2^F/F^* innate immune cells are impaired in pathogenic clearance functions. Whole bone marrow was harvested from vehicle- or MPLA-pretreated mice infected with *P. aeruginosa*, and ROS was assessed via flow cytometry in **(A)** all cells, **(B)** macrophages, and **(C)** neutrophils. Peritoneal fluid was similarly collected and assessed for ROS production in **(D)** macrophages and **(E)** neutrophils. **(F)** Schematic of *in vivo* phagocytosis assessment. Mice were pretreated with MPLA or vehicle for 24 hours prior to injection with pHrodo beads, and phagocytosis was assessed 6 hours later in **(G)** macrophages (F4/80^+^), cell number positive for pHrodo, and **(H)** geometric MFI (gMFI) of pHrodo, as well as **(I)** neutrophils (Ly6G^+^) as cell number positive for pHrodo. Data are shown as mean ± SEM, two-way ANOVA with post-hoc Tukey’s test, n = 15-28 mice per group for peritoneal ROS, n = 15-23 mice per group for BM ROS, and n = 13-17 mice per group for phagocytosis. Each point represents an individual mouse.

### *VavTet2^F/F^* mice display impaired myeloid differentiation during infection and MPLA pretreatment

We then asked whether these deficits in innate immune cell function are reflected in differentiation. We characterized BM innate immune cell populations by spectral flow cytometry from LR- or MPLA-pretreated mice infected with *P. aeruginosa* (gating strategy in Supplemental Figure 4). Total counts and frequencies of mature neutrophils (CD11b^+^ Ly6G^high^) increase similarly when administered MPLA before infection (Figure 5A-B). We confirmed that no changes were due to baseline or MPLA-induced effects (Supplementary Figure 5A-B). Interestingly, *VavTet2^F/F^* mice increase less-mature BM neutrophils (CD11b^+^ Ly6G^mid^) relative to WT and *VavTet2^F/F^Ripk1^D/+^* during infection (Figure 5C-D). Similar to neutrophils, infected *VavTet2^F/F^* mice augment macrophage production (CD11b^+^ F4/80^+^), but deplete mature macrophages (CD11b^+^ F4/80^high^) in a RIPK1-dependent manner with MPLA prior to *P. aeruginosa* (Figure 5E-H, Supplemental Figure 5E-H). Relative accumulation of immature neutrophils is consistent with impaired differentiation in *VavTet2^F/F^* mice that can be improved with *RIPK1* kinase genetic inactivation.

**Figure 5:**
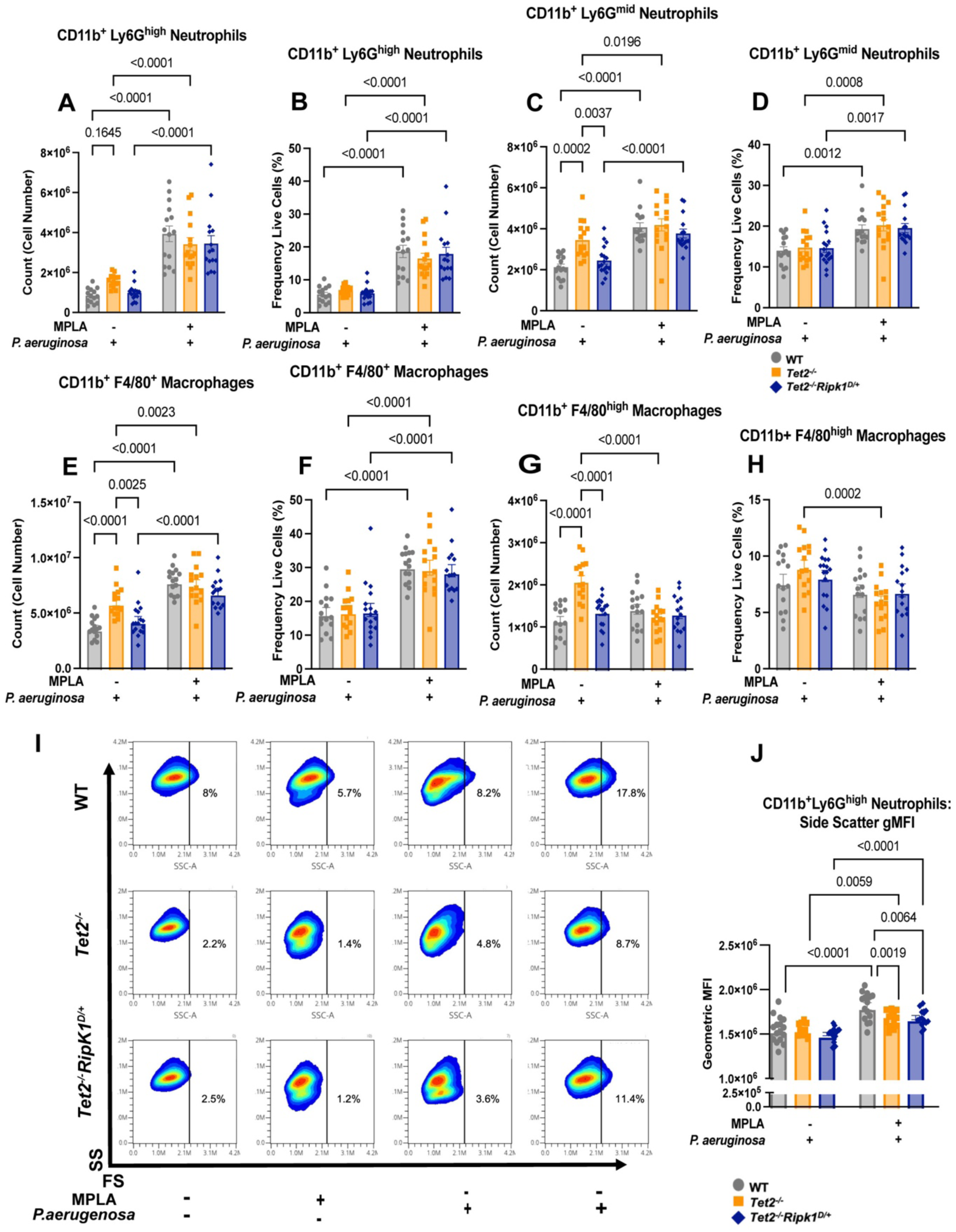
*VavTet2^F/F^*mice display impaired myeloid differentiation during infection and MPLA pretreatment. Whole bone marrow from vehicle- or MPLA-pretreated and *P. aeruginosa* infected mice was assessed via flow cytometry for mature innate immune cell populations. **(A-B)** Mature neutrophils (CD11b^+^ Ly6G^high^), **(C-D)** less mature neutrophils (CD11b^+^, Ly6G^mid^) **(E-F)** macrophages (CD11b^+^ F4/80^+^) and **(G-H)** mature macrophages (CD11b^+^ F4/80^high^) counts and frequencies were assessed. **(I)** Combined flow plots from all treatment groups (LR- and MPLA-pretreated, and mock or *P. aeruginosa* infected). Gated mature neutrophils (CD11b^+^ Ly6G^high^) were plotted for FSC and SSC, and frequencies as FSC high are shown. **(J)** Quantified geometric mean fluorescent intensity (gMFI) of SSC for vehicle- and MPLA-pretreated and *P. aeruginosa* infected groups within mature neutrophils. Data are shown as mean ± SEM, two-way ANOVA with post-hoc Tukey’s test, n = 15-17 mice per group. Each point represents an individual mouse.

Given impaired neutrophil differentiation, we examined mature neutrophil size and granularity. *VavTet2^F/F^* and *VavTet2^F/F^Ripk1^D/+^*mice decrease FSC, suggesting smaller size, across all treatment groups (Figure 5I). Similarly, *VavTet2^F/F^* and *VavTet2^F/F^Ripk1^D/+^* mature neutrophils are less complex in MPLA- pretreated and infected mice relative to WT, quantified by geometric MFI (gMFI) of side scatter (Figure 5J, Supplemental Figure 6A). Immature neutrophils are similar in size and complexity across genotypes (Supplemental Figure 6B-C). Finally, no statistical differences emerge across genotypes in CD62L expression, though similarly, *Tet2^-/-^*mice have trends towards higher CD62L^mid^ Ly6G^+^ neutrophil counts (Supplemental Figure 7A-J). Collectively, these data indicate that infection-initiated inflammatory states promote differentiation deficits in neutrophils during *Tet2* loss. RIPK1 kinase inactivation improves neutrophil differentiation measured by surface markers, bit size and granularity remain altered.

Lastly, *VavTet2^F/F^* but not WT or *VavTet2^F/F^Ripk1^D/+^* mice increase dendritic cell (DC; CD11b^+^ CD11c^+^) counts with infection, and MPLA prior to infection decreases BM DCs in *VavTet2^F/F^*mice in a RIPK1- dependent manner (Supplemental Figure 8A-B). Ter119^+^ cells decrease in *VavTet2^F/F^*mice during infection (Supplemental Figure 8C-D). Finally, we show no differences between genotypes in B cells, suggesting limited adaptive immune system involvement in this model (Supplemental Figure 8E-F).

### MPLA with *P. aeruginosa* promotes myeloid bias at the expense of erythroid and megakaryocytes in

#### VavTet2^F/F^ mice

*Tet2* loss directs myeloid-biased hematopoiesis following TLR4 agonist challenge, in aging, and CH.^12^ Additionally, myeloid-biased hematopoiesis is a key feature of infection response. To determine whether infection exaggerates myeloid-biased hematopoiesis and whether MPLA prior to infection exacerbates myeloid-bias in *VavTet2^F/F^* mice, we evaluated hematopoietic stem and progenitor populations by spectral flow cytometry (gating strategy in Supplemental Figure 9).^41^

We first evaluated our data in an unbiased manner to identify statistically significant stratifying signatures between genotypes for each treatment group. Cluster identification, characterization, and regression (CITRUS) analysis revealed that at baseline, the lineage negative (Lin^-^) population was the stratifying signature between genotypes. In all other conditions, (MPLA, infection, and MPLA + infection), Sca-1 was the stratifying marker, demonstrating in an unbiased manner that surface Sca-1 is significantly increased in *VavTet2^F/F^* mice upon inflammatory challenge (data not shown).

Sca-1 expression changes due to inflammatory states, and *Tet2* loss also increases this in response to inflammation.^12,42,43^ An alternative gating strategy has been proposed using CD86 as a reliable substitute for Sca-1 to denote LSK populations, thereafter referred to as “L86K.”^44^ We therefore compared the behavior of these two methods in this innate immune memory and infection model by characterizing hematopoietic populations in WT, *VavTet2^F/F^*, and *VavTet2^F/F^Ripk1^D/+^* mice.^44^ *VavTet2^F/F^* but not WT or *VavTet2^F/F^Ripk1^D/+^* BM significantly increase LSK populations relative to L86K with infection and MPLA prior to infection (Figure 6A). We also examined this effect in Lin^-^ populations via t-distributed Stochastic Neighbor Embedding (tSNE) plots to visualize upregulations in Sca-1 expression with inflammatory stimulation. Both at baseline and with *P. aeruginosa*, Sca-1 expression increases in *VavTet2^F/F^* mice compared to WT, and infection exacerbates this effect (Figure 6B, Supplemental Figure 10A). However, CD86 expression remains consistent regardless of genotype or inflammatory stimulation (Figure 6B, Supplemental Figure 10B). Therefore, inflammation may artifactually enhance LSK populations in *VavTet2^F/F^* hematopoietic stem and progenitor populations, and we moved forward using L86K gating, although we report Sca-1-based gating in the Supplemental Information for comparison.

**Figure 6:**
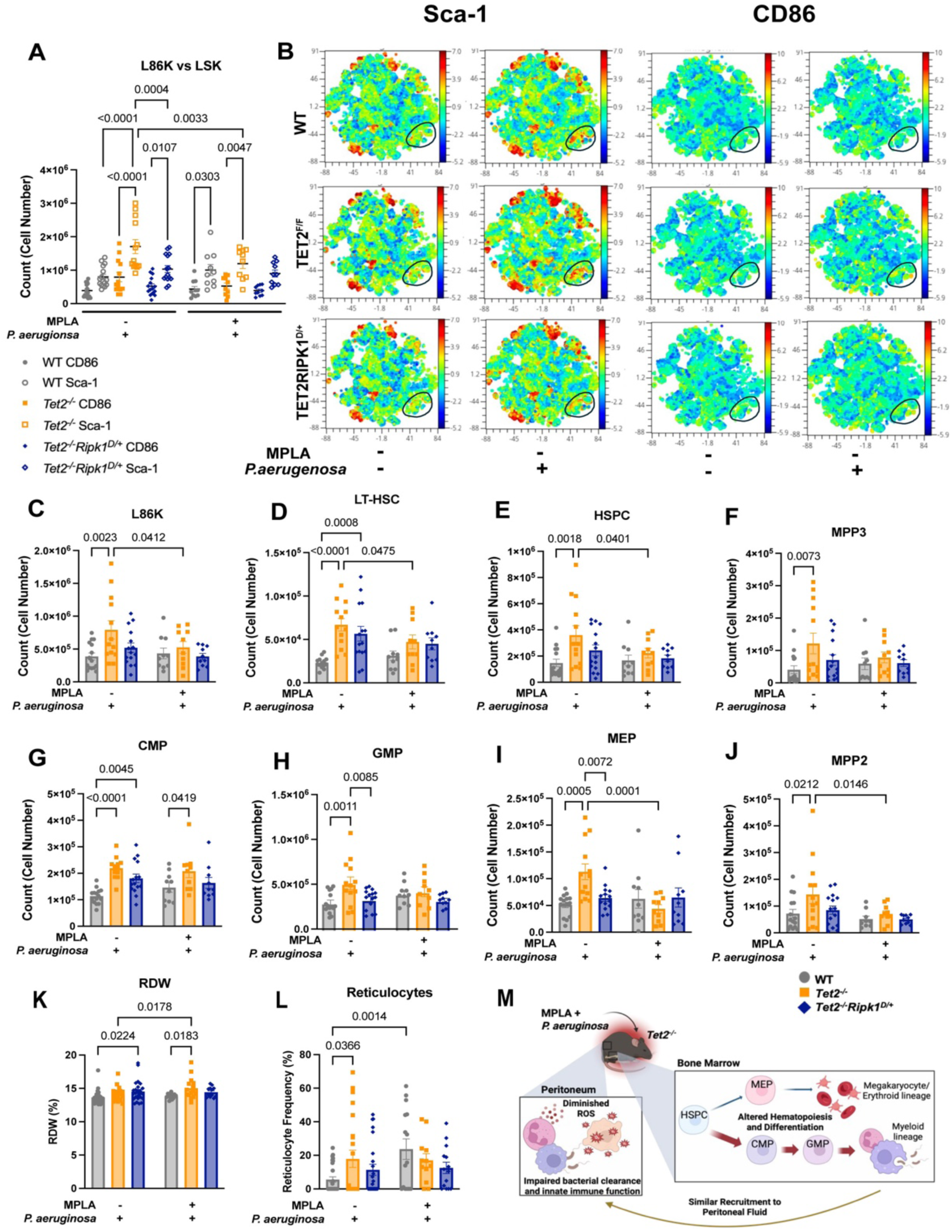
MPLA with *P. aeruginosa* promotes myeloid bias at the expense of erythroid and megakaryocytes in *VavTet2^F/F^*mice. Whole bone marrow from vehicle- and MPLA-pretreated mice was harvested for spectral flow cytometry of bone marrow stem and progenitors. **(A)** Comparison of CD86 and Sca- 1 gating. **(B)** tSNE comparison plots of Sca-1 (left) and CD86 expression (right) within Lin^-^ populations gated from live cells, all biological replicates overlaid on each plot, n = 5-16 biological replicates. Total cell counts are shown for **(C)** L86K stem and progenitors, and via CD86 gating, **(D)** long-term hematopoietic stem cells (LT-HSC), **(E)** hematopoietic stem and progenitors (HSPC), **(F)** multipotent progenitor subset 3 (MPP3), **(G)** common myeloid progenitor (CMP), **(H)** granulocyte-monocyte progenitor (GMP), **(I)** megakaryocyte- erythroid progenitor (MEP), and **(J)** multipotent progenitor subset 2 (MPP2). Whole blood was collected from mice, and CBC performed. Shown are **(K)** red-cell distribution width (RDW), and **(N)** reticulocyte frequency. **(M)** Summary of results. Data are shown as mean ± SEM, two-way ANOVA with post-hoc Tukey’s test, n = 10- 16 mice per group for flow cytometric data and n = 15-28 mice for CBC data. Each point represents an individual mouse.

We then examined hematopoietic perturbations induced by MPLA with infection in hematopoietic stem and intermediate progenitors. Intriguingly, infection expands L86K, hematopoietic stem and progenitor cell (HSPC), and long-term hematopoietic stem cell (LTHSC) populations in *VavTet2^F/F^* BM relative to WT BM, dampened by RIPK1 inactivation (Figure 6C-E). While the size of WT and *VavTet2^F/F^Ripk1^D/+^* L86K, HSPC, and LTHSC populations is not influenced by MPLA prior to *P. aeruginosa*, these populations decrease in *VavTet2^F/F^* BM. RIPK1 kinase inactivation (*VavTet2^F/F-^Ripk1^D/+^* BM) prevents exaggerated expansion of L86K, HSPC, and LTHSC populations following infection and preserves these populations in MPLA-pretreated and infected mice (Figure 6C-E). Exaggerated expansion in *VavTet2^F/F^* L86K, HSPC, and LTHSC populations is not due to baseline differences or MPLA alone and occurs in short-term hematopoietic stem cells (ST-HSC; Supplementary Figure 11A-D). Next, we investigated myeloid lineage-biased progenitors, including multipotent progenitor subset 3 (MPP3), common myeloid progenitor (CMP), and granulocyte-monocyte progenitor (GMP) (Figure 6F-H, Supplemental Figure 11E-G). We observe a similar effect in infected *VavTet2^F/F^* mice; these populations expand relative to WT, though RIPK1 inactivation partially dampens this effect. MPLA with infection does not deplete these populations, indicating preferential expansion of myeloid-biased progenitors during inflammation.

To determine if myeloid-biased expansion occurs at the expense of other lineage-biased stem and progenitor populations, we examined erythroid/megakaryocyte-biased progenitors. Though infection expands multipotent progenitor population 2 (MPP2) and megakaryocyte-erythroid progenitors (MEP), MPLA pretreatment depletes these subsets, indicating relative reduction of the megakaryocyte/erythroid-biased hematopoietic compartment (Figure 6I-J, Supplemental Figure 11H-I). A similar trend occurs in multipotent progenitor population 4 (MPP4), which are lymphoid lineage-biased (Supplemental Figure 11J), suggesting that additional lineages may be affected by preferential myeloid expansion. Similar trends occur in frequency of live cells (Supplemental Figure 12A-J). Additionally, employing Sca-1 gating strategies in place of CD86 reveals the same trends, though effects are much greater, and differences occur in stem and progenitors between genotypes at baseline (Supplemental Figure 13A-N).

Our experimental paradigm induces sepsis and is terminated six hours post-infection. Even within this short timeframe, *VavTet2^F/F^* mice increase peripheral blood red cell distribution width (RDW), an indicator for immature erythrocyte production and stress erythropoiesis with MPLA prior to *P. aeruginosa* (Figure 6K). This effect is corroborated by increased reticulocytes in MPLA-pretreated and infected *VavTet2^F/F^* mice, demonstrating persistent stress erythropoiesis (Figure 6L).

Taken together, these results reveal that *VavTet2^F/F^* mice fail to generate effective innate immune responses to *P. aeruginosa* (Figure 6M). Initiation of innate immune memory by MPLA pre-treatment improves subsequent response to *P. aeruginosa,* however this response is diminished relative to WT. This may be due to impaired myeloid differentiation, as *VavTet2^F/F^* myeloid cells exhibit defective differentiation, altered granularity, and size (Figure 5). Importantly, MPLA prior to *P. aeruginosa* expands myeloid-biased progenitors at the expense of megakaryocyte/erythroid-progenitors in *VavTet2^F/F^* mice, which is improved by RIPK1 inactivation.

## DISCUSSION

Inflammatory states promote *Tet2*-mutant cell expansion, which can then promote hyperinflammatory responses to these stimuli, inducing a feed-forward inflammatory process that further impairs hematopoiesis.^11–14^ Additionally, TET2 mutations impair innate immune responses, impeding infection clearance, phagocytosis, and others.^9,10^ Inability to clear infections allows for inflammatory persistence, creating the selective niche for *Tet2*-mutant cell expansion and disease progression. RIPK1 is furthermore involved in inflammatory signaling pathways that may promote hyperinflammatory response of *Tet2* loss.^39,40,45^ MPLA-induced innate immune memory additionally boosts innate immune responses to secondary infection while simultaneously dampening proinflammatory responses.^17,19,23,28,29^

We show that *Tet2*-deficiency upregulates inflammation during LPS in a RIPK1-dependent manner within BM. IL-1β and TNF-α are critical for *Tet2*-mediated disease progression, and TNF-α also functions in HSC survival and myeloid expansion.^13,14,32,46^ IL-6 mediates expansion of *Tet2^-/-^* hematopoietic stem and progenitor cells following an inflammatory stimulus with LPS through the SHP2-Stat3 signaling axis.^12^ In other models of *Tet2* loss, IL-6 is variably altered in a context and model dependent manner,^11,12,47,49,50^ indicating that environmental and context-specific effects can sway the impact of *Tet2* loss on inflammatory cytokines.^12^

Our finding that serum cytokines are not altered in *Tet2* loss relative to WT during infection are consistent with other reported infectious models (*Escherichia* coli and *P. aeruginosa*) ^30,47,48^ *VavTet2^F/F^* mice maintain serum cytokine levels comparable to WT counterparts. Though augmented inflammatory response to infection may be anticipated in *Tet2* loss, changes in circulating cytokines are dynamic and not captured in all models. Alternatively, enhanced cytokine production may occur within the BM microenvironment or in an autocrine manner. Direct sensing of cytokines via TLRs on HSCs could induce hematopoietic perturbations observed in this model.^30^

*Tet2* loss in mature myeloid cells via *LysMCre* upregulates IL-6 and IL1-β in macrophages following *ex vivo* stimulation with *E. coli* or *P. aeruginosa.*^47,48^ Additionally, *LysMCreTet2^fl/fl^*mice upregulate IL-6 *in vivo* following *E. coli* or *P. aeruginosa*.^47,48^ This contrasts to our findings, which show no difference in IL-6 or other serum cytokines in *VavCreTet2^fl/fl^* compared to WT following *P. aeruginosa* (Figure 1). These results highlight the importance of developmental stage of TET2 loss and infectious agent on inflammatory response to infectious insult.

Several ground-breaking studies probed the impact of infection and autoimmunity on HSCs.^15,25,49–51^ Our study explores whether innate immune memory may be harnessed to improve infection response in *VavTet2^fl/fl^* mice. Impaired bacterial clearance and innate immune functions are known features of *Tet2*- deficiency,^9,10,54^ We find similarly that *VavTet2^F/F^* mice are impaired in response to *P. aeruginosa*. MPLA prior to *P. aeruginosa* markedly improves infection response in *VavTet2^F/F^* mice, indicating that *Tet2* loss does not prevent innate immune memory. However, *Tet2* loss drives functional and differentiation impairments in innate immune cells that reduce bacterial clearance capacity relative to WT even after MPLA pre-treatment.

Although MPLA improves infection response in *VavTet2^F/F^* mice, an aberrant hematopoietic stem and progenitor cell response occurs.^55^ *VavTet2^F/F^* mice display exaggerated myeloid bias, a key characteristic of MDS, at the expense of erythroid and megakaryocyte depletion at the stem and progenitor cell level.

Interestingly, RIPK1 kinase inactivation does not alter response to infection or innate immune memory but markedly affects aberrant hematopoietic stem and progenitor cell responses driven by *Tet2* loss. These results suggest a role for hematopoietic stem and progenitor cells in sensing inflammation in *Tet2*-deficiency, demonstrating an exaggerated and aberrant response with myeloid skewing, relative erythroid depletion, and aberrant myeloid differentiation that is dampened by RIPK1 kinase inhibition. Moreover, this suggests that MPLA with infection initiates thrombocytopenias and anemias, clinical markers of MDS, at the stem cell level during *Tet2* loss which is dampened with RIPK1 inactivation. This warrants further investigation into RIPK1 kinase inhibition to improve hematopoietic dysfunction during inflammation.^56^

Although 6-hour infection captures acute Gram-negative bacterial response, additional studies are necessary to determine whether innate immune memory protects *Tet2*-deficient organisms in longer infection models. Moreover, it is important to determine if MPLA cross-protects *Tet2^-/-^* mice against Gram-positive or other infections.

Collectively, this study shows that MPLA prior to infection in *Tet2* loss prevents severe bacterial infection, yet MPLA treatment promotes MDS-like disease in a RIPK1-dependent manner. Our results provide critical insight into the effects of infection-mediated inflammation on *Tet2*-driven disease, which warrant additional investigation, as the findings hold therapeutic potential for improving infection response and preventing disease progression.

## ACKNOWLEDGEMENTS

This work was supported by grants from the National Heart, Lung, and Blood Institute (R01HL133559) (S.Z.), Department of Veteran’s Administration (I01BX002250, I01BX004365 (renewal) (S.Z.), National institute of General Medical Sciences, (5R35GM141927) (J.K.B.), and National Institute of Allergy and Infectious Diseases (5R01AI151210) (E.R.S.). We thank Michelle Kelliher for RIPK1^D138N^ mice and Omar Abdel-Wahab for TET2^fl/fl^ mice. We thank Christian Warren of the VA Tennessee Valley healthcare system flow cytometry facility for assistance and feedback with spectral flow cytometry. We thank Edward R. Sherwood for thoughtful discussion and feedback and for providing resources. We thank Yuliya Hassan for contributions to animal care and genotyping. We thank Sophia Cornish for contributions to qRT-PCR in Supplementary Figure 1. Schematics created in the graphical abstract and in Figures 2, 4, and 6 created using BioRender.com. A.N.L.J. is a Ph.D. candidate at Vanderbilt University and this work is submitted in partial fulfillment of the requirement for the Ph.D. The content of this article contains opinions solely of the authors, and they do not necessarily reflect the views or policies of the Department of Veteran’s Administration, the US government, or the National Institutes of Health.

## AUTHOR CONTRIBUTIONS

Contributions: A.N.L.J., X.D., M.O., Y.W., J.K.B., and S.Z. performed experiments. A.N.L.J., X.D., M.O., H.C., J.K.B., and S.Z. analyzed results. A.N.L.J., Y.W., H.C., and S.Z. made the figures. A.N.L.J. and S.Z. wrote the paper, and A.N.L.J., X.D., M.O., J.K.B., and S.Z. edited the paper. A.N.L.J., X.D., J.K.B., and S.Z. conceptualized this project.

## SUPPLEMENTAL METHODS

### Mice

*Tet2^fl/f^* mice and *Ripk1^D^*^138^*^N/D^*^138^*^N^* mice were obtained from Omar Abdel-Wahab and Michelle Kelliher, respectively.^1,2^ *vavCreTet2^fl/+^* mice were generated by crossing *vavCre+* mice with *Tet2^fl/fl^* mice, which were backcrossed with *Tet2^fl/fl^* mice to generate *vavCreTet2^fl/fl^* mice (hereafter *Tet2^-/-^*). *Ripk1^D^*^138^*^N/+^* mice were crossed with *vavCre+* mice to generate *vavCreRipk^D^*^138^*^N/+^*mice. These were crossed to *Tet2^fl/fl^* mice to generate *vavCreTet2^fl/+^Ripk1^D^*^138^*^N/+^* and were backcrossed with *Tet2^fl/fl^* mice to generate the *vavCreTet2^fl/fl^Ripk1^D^*^138^*^N/+^* mice (hereafter *Tet2^-/-^Ripk1^D/+^*). Mice were backcrossed at least 9 generations with C57Bl/6J mice from The Jackson Laboratory. Both male and female mice, aged 6-34 weeks, were used for experiments. Mice were age- and sex-matched across genotypes and treatment groups.

LPS treatment of mice was performed by intraperitoneally (IP) injecting mice with 1.5mg/kg LPS (L4391; Millipore-Sigma, Burlington, MA). MPLA (vac-mpla2, Invivogen, San Diego, CA) was diluted in LR to 100 μg/mL and injected into mice intraperitoneally at 20 μg in 0.2 mL.

### Infection of mice

*Pseudomonas aeruginosa* (Schroeter) Migula (ATCC 19660, Manassas, Virginia) was thawed, and entire vial contents were cultured for 22 hours in tryptic soy agar (TSA) broth in incubator. *P. aeruginosa* were then diluted to 1×10^8^ CFU/mL. Mice were sprayed with ethanol, and 1 mL was injected intraperitoneally per mouse.

### Bone Marrow Derived Macrophage Culture

To generate L-929 conditioned media, L929 cells (NCTC clone 929 CCL-1, ATCC, Manassas, Virginia) were cultured in DMEM with 4.5 g/dL glucose (10017CV, Corning Inc., Corning, NY), L-Glutamine (25-005-CI, Corning Inc., Corning, NY), 25mM HEPES (H3662 Millipore-Sigma, Burlington, MA), 10% FBS (10437-028, ThermoFisher Scientific, Waltham, MA), and 1% Penn/Strep (15140-122, ThermoFisher Scientific, Waltham, MA) for 7 days, adding additional DMEM on day 3. GM-CSF conditioned media was collected from the cell monolayer and filtered using a 0.25 µm filter.

Mouse femur and tibia were harvested from mice, and bone marrow was harvested as previously described.^3^ All bone marrow was then plated in a 100mm petri dish and cultured in DMEM with 10% FBS, 2 mM L-Glutamine, and 1% Penn/Strep with 10% L929-conditioned media for 7 days. BMDMs were then removed from the dish via trypsinization (0.25% Trypsin EDTA, 25200-056, ThermoFisher Scientific, Waltham, MA) and plated on a 60mm dish in DMEM10 with 10% L929-conditioned media until downstream use.

### Serum cytokines

Plasma was obtained from mice, and serum cytokine concentration was determined via Bio-Rad Bio-Plex Pro Mouse Cytokine Assay according to manufacturer guidelines, complete list of reagents in Supplemental Table 1. Imaging was performed on a Bio-Plex Magpix Multiplex Reader (Bio-Rad, Hercules, CA).

### RNA Isolation and Quantitative Reverse-Transcriptase PCR

RNA was isolated and purified from BMDMs using the RNeasy Mini Kit (74104, Qiagen, Hilden, Germany) or Quick-RNA MiniPrep (R1054, Zymo Research, Irvine, CA) according to manufacturer protocols. Concentration and purity of RNA was determined via NanoDrop One (ThermoFisher Scientific, Waltham, MA). cDNA was synthesized from RNA with the SuperScript™ III First-Strand Synthesis System (18080051, ThermoFisher Scientific, Waltham, MA) according to manufacturer guidelines. qPCR was then performed using the TaqMan™ Gene Expression Assay (4331182, Mm00434228_m1 Il1b, Mm00446190_m1 Il6, Mm00443258_m1 Tnf, Mm02619580_g1 Actb, ThermoFisher Scientific, Waltham, MA) and run using Bio-rad C1000 Touch Thermal Cycler CFX96 Real-Time System (Bio-Rad, Hercules, CA). Data were analyzed using the ddCT method.^4^ Experimental samples were run in technical duplicates, then CT values were averaged for each duplicate. β-actin was used as a housekeeping control. Data were normalized to PBS within genotypes.

### Bacterial Burden Assessments

Peritoneal cavity was aseptically flushed with 5 mL of cold PBS. Serial dilutions were performed from peritoneal lavage, and 100 µL were cultured overnight on TSA and colony forming units were quantified.

### Reactive Oxygen Species Production

Respiratory burst buffer (RBB; 1 g/mL BSA, 1 µM CaCl_2_) and 1× DHR were prepared from kit (601130, Cayman Chemicals, Ann Arbor, MI). 1×10^6^ cells were added to flow tubes and washed with RBB, incubated, then stained with DHR for 1 hour. Cells were washed with 2 mL PBS and resuspended in RBB prior to assessment via BD Accuri C6 Plus Flow Cytometer (BD BioSciences, Franklin Lakes, NJ). Monocytes, neutrophils, and macrophages were gated via forward and side scatter populations.

### Phagocytosis Assay

Mice were pretreated with MPLA or vehicle control (as above) 1 day prior to assessment. pHrodo™ Red E. coli BioParticles™ Conjugate for Phagocytosis (P35361, Invitrogen, San Diego, CA) were prepared by resuspending in sterile saline and sonicating for 5 minutes at room temperature and intraperitoneally injected into mice. Six hours later, peritoneal cells were harvested by washing the peritoneum with 5 mL cold PBS, and cells were stained with fluorochrome-conjugated antibodies (Supplemental Table 2) prior to phagocytosis assessment via BD Accuri C6 Plus Flow Cytometer (BD BioSciences, Franklin Lakes, NJ).

### Flow Cytometry of Infected Murine Bone Marrow and Peritoneal Cells

Bone marrow was harvested as above and passed through a 70 µm filter. Red blood cells were lysed (15 mM NH_4_Cl, 12 µM NaHCO_3_, 1 mM EDTA) and passed through a 70 µm filter. Peritoneal cells were collected by flushing the peritoneum with 5 mL PBS and passing through a 70 µm filter. 2 x 10^6^ cells were added to a 96- well round-bottom plate and stained for viability. Cells were pelleted and incubated with fluorochrome-conjugated antibodies (Supplemental Table 3). Cells were then fixed in 2% paraformaldehyde until assessment via BD Cytek® Aurora (Cytek® Biosciences, Fremont, California).

### Spearman correlation, PCA, and Nanostring analysis

NanoString nCounter Mouse Myeloid Innate Immunity V2 Panel was used, which contains 754 genes, including 20 internal reference genes for data normalization to provide comprehensive coverage of inflammatory and immune-related signaling. Log2 FC (Log2 fold change) was calculated as the difference in gene expression (log2 scale) between sample groups. P value was adjusted (adjusted P value) for multiple comparisons based on False Discovery Rate (FDR) method following Limma t test to find differentially expressed genes between sample groups (raw P value). Spearman correlation heatmap generated and clustered based on gene expression in log2 scale. PCA plot Principal component analysis (PCA) generated by Prcomp package in R based on log2 transformed gene expression.

## SUPPLEMENTAL TABLES

**Supplemental Table 1:** List of reagents used for Bio-Plex Cytokine assays.

| Reagent | Vendor | Catalog Number |
| --- | --- | --- |
| Bio-Plex Pro Mouse Cytokine Assay | Bio-Rad | 12022433 and M60000007A |
| Bio-Plex Pro™ Mouse Cytokine MIP-2 Set | Bio-Rad | 171G6006M |
| Bio-Plex Pro™ Mouse Cytokine G-CSF Set | Bio-Rad | 171G5015M |
| Bio-Plex Pro Mouse Cytokine IL-6 Set | Bio-Rad | 171G5007M |
| Bio-Plex Pro Mouse Cytokine Group II Standards, 96 assays | Bio-Rad | 171I60001 |

**Supplemental Table 2:** List of antibodies used in *in vivo* phagocytosis assays.

| Reagent | Vendor | Catalog Number | Concentration (mg/mL) |
| --- | --- | --- | --- |
| BD Pharmingen™ FITC Rat anti-Mouse Ly-6G | BD Biosciences | 551460 | 0.5 |
| F4/80 Monoclonal Antibody (BM8), FITC, eBioscience™ | Invitrogen | 11-4801-82 | 0.5 |
| Ly-6C Monoclonal Antibody (HK1.4), PerCP-Cyanine5.5, eBioscience™ | Invitrogen | 45-5932-82 | 0.2 |
| BD Pharmingen™ Purified Rat Anti-Mouse CD16/CD32 (Mouse BD Fc Block™) | BD Biosciences | 553142 |  |
| Rat IgG2a kappa Isotype Control (eBR2a), FITC, eBioscience™ | Invitrogen | 11-4321-80 | 0.5 |
| Armenian Hamster IgG Isotype Control(eBio299Arm), PerCP-Cyanine5.5, eBioscience™ | Invitrogen | 45-4888-80 | 0.2 |
| Rat IgG2b kappa Isotype Control (eB149/10H5), PE, eBioscience™ | Invitrogen | 12-4031-82 | 0.2 |

**Supplemental Table 3:** List of antibodies used for spectral flow cytometry assays.

| Reagent | Vendor | Catalog Number | Concentration (mg/mL) |
| --- | --- | --- | --- |
| LIVE/DEAD fixable Aqua Dead Cell Stain Kit | ThermoFisher Scientific | L34965 | 1:1000 |
| BD Horizon™ BV605 Rat Anti-Mouse CD127 | BD Biosciences | 569295 | $1 \times 10^{-3}$ |
| BD Horizon™ BV650 Rat Anti-Mouse CD62L | BD Biosciences | 564108 | $2.5 \times 10^{-4}$ |
| Ly-6A/E (Sca-1) Monoclonal Antibody (D7), Brilliant Violet™ 786, eBioscience™ | ThermoFisher Scientific | 417-5981-80 | $1 \times 10^{-3}$ |
| CD48 Monoclonal Antibody (HM48-1), FITC, eBioscience™ | ThermoFisher Scientific | 11-0481-81 | $3.125 \times 10^{-4}$ |
| CD184 (CXCR4) Monoclonal Antibody (2B11), PerCP-eFluor™ 710, eBioscience™ | ThermoFisher Scientific | 46-9991-80 | $5 \times 10^{-4}$ |
| CD135 (Flt3) Monoclonal Antibody (A2F10), PE-Cyanine5, eBioscience™ | ThermoFisher Scientific | 15-1351-81 | $1 \times 10^{-3}$ |
| CD150 Monoclonal Antibody (mShad150), PE-Cyanine7, eBioscience™ | ThermoFisher Scientific | 25-1502-80 | $8.33 \times 10^{-5}$ |
| BD Pharmingen™ Alexa Fluor® 647 Rat anti-Mouse CD34 | BD Biosciences | 560230 | $4 \times 10^{-3}$ |
| BD Pharmingen™ APC-H7 Rat anti-Mouse CD117 | BD Biosciences | 560185 | $1.25 \times 10^{-4}$ |
| CD71 (Transferrin Receptor) Monoclonal Antibody (R17217 (RI7 217.1.4)), eFluor™ 450, eBioscience™ | ThermoFisher Scientific | 48-0711-82 | $1.25 \times 10^{-4}$ |
| FITC Anti-Mouse F4/80 Antigen (BM8.1) | Tonbo Biosciences | 35-4801 | $6.25 \times 10^{-4}$ |
| BD Pharmingen™ PE Rat Anti-Mouse CD41 | BD Biosciences | 558040 | $1.25 \times 10^{-4}$ |
| BD Horizon™ PE-CF594 Rat Anti-CD11b | BD Biosciences | 562317 | $6.25 \times 10^{-5}$ |
| BD Pharmingen™ PE-Cy™5 Rat Anti-Mouse CD45R/B220 | BD Biosciences | 553091 | $1.25 \times 10^{-4}$ |
| BD Pharmingen™ PE-Cy™7 Rat Anti-Mouse Ly-6G | BD Biosciences | 560601 | $6.25 \times 10^{-5}$ |
| APC Anti-Mouse CD11c (N418) | Tonbo Biosciences | 20-0114 | $1 \times 10^{-3}$ |
| TER-119/Erythroid Cells Rat anti-Mouse, Alexa Fluor 700, Clone: TER-119 | BD Biosciences | 560508 | $2.5 \times 10^{-4}$ |
| F4/80 Monoclonal Antibody (BM8), eFluor™ 450, eBioscience™ | ThermoFisher Scientific | 48-4801-82 | $5 \times 10^{-4}$ |
| BD Pharmingen™ Biotin Hamster Anti-Mouse CD3e | BD Biosciences | 553060 | $1 \times 10^{-3}$ |
| BD Pharmingen™ Biotin Rat Anti-Mouse Ly-6G and Ly6C | BD Biosciences | 553125 | $1 \times 10^{-3}$ |
| BD Pharmingen™ Biotin Rat Anti-Mouse TER-119/Erythroid Cells | BD Biosciences | 553672 | $1 \times 10^{-3}$ |
| BD Pharmingen™ Biotin Rat Anti-Mouse CD45R/B220 | BD Biosciences | 553086 | $1 \times 10^{-3}$ |
| BD Pharmingen™ Purified Rat Anti-Mouse CD16/CD32 (Mouse BD Fc Block™) | BD Biosciences | 553142 | $2.08 \times 10^{-4}$ |
| CD86 Rat anti-Mouse, Alexa Fluor 700, Clone:GL1 | Fisher Scientific | BDB560581 | $2 \times 10^{-3}$ |
| eBioscience™ Streptavidin eFluor™ 450 Conjugate | eBioscience | 48-4317-82 | $8.33 \times 10^{-5}$ |

## SUPPLEMENTAL FIGURES

**Supplemental Figure 1:**
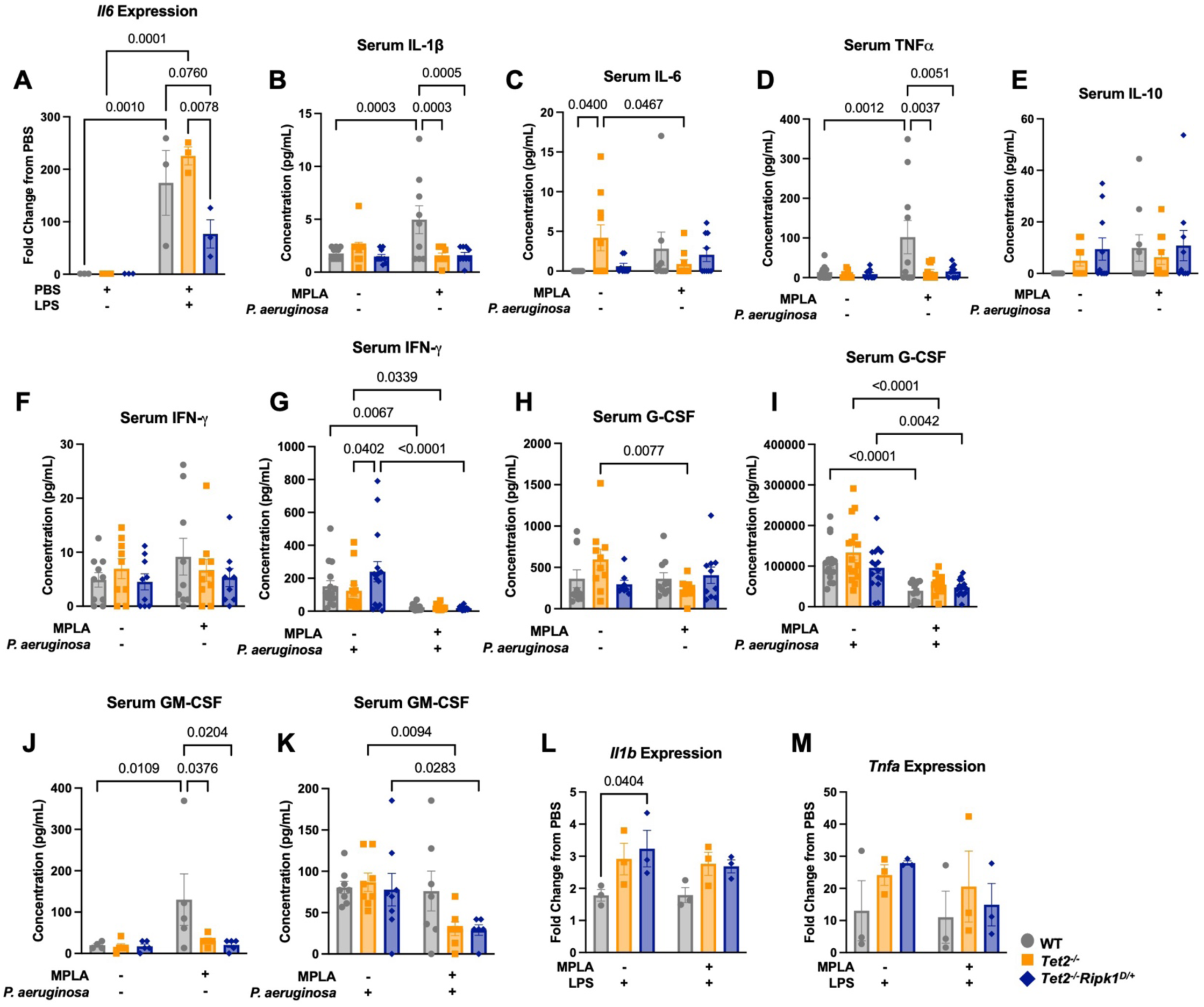
Cytokine profile of *in vivo* and *in vitro* inflammatory challenge. Mice were treated injected with LPS intraperitoneally, and whole bone marrow was harvested 8 hours later. qRT-PCR was performed for **(A)** *Il6.* Data were normalized to PBS control within each genotype. Mice were given vehicle or MPLA for 3 days, following which they were either mock infected or infected with *P. aeruginosa* or vehicle for 6 hours. Following infection, whole blood was collected, and plasma was isolated. A Bio-Rad Bio-Plex assay was performed to determine concentration of circulating serum cytokines. Shown are **(B)** IL-1β, **(C)** IL-6, **(D)** TNF-α, **(E)** IL-10, **(F-G)** IFN-γ, **(H-I)** G-CSF, and **(J-K)** GM-CSF. BMDMs were stimulated with PBS or MPLA for 3 days, and LPS was given for 24 hours. Expression of proinflammatory genes was quantified by qRT-PCR and shown are **(L)** *Il1b* and **(M)** *Tnfa* expression. Data were normalized to PBS control within genotype. n = 3 biological replicates for qRT-PCR, n = 5-16 biological replicates for Bioplex. Data are shown as mean ± SEM, 2-way ANOVA with post-hoc Tukey’s test.

**Supplemental Figure 2:**
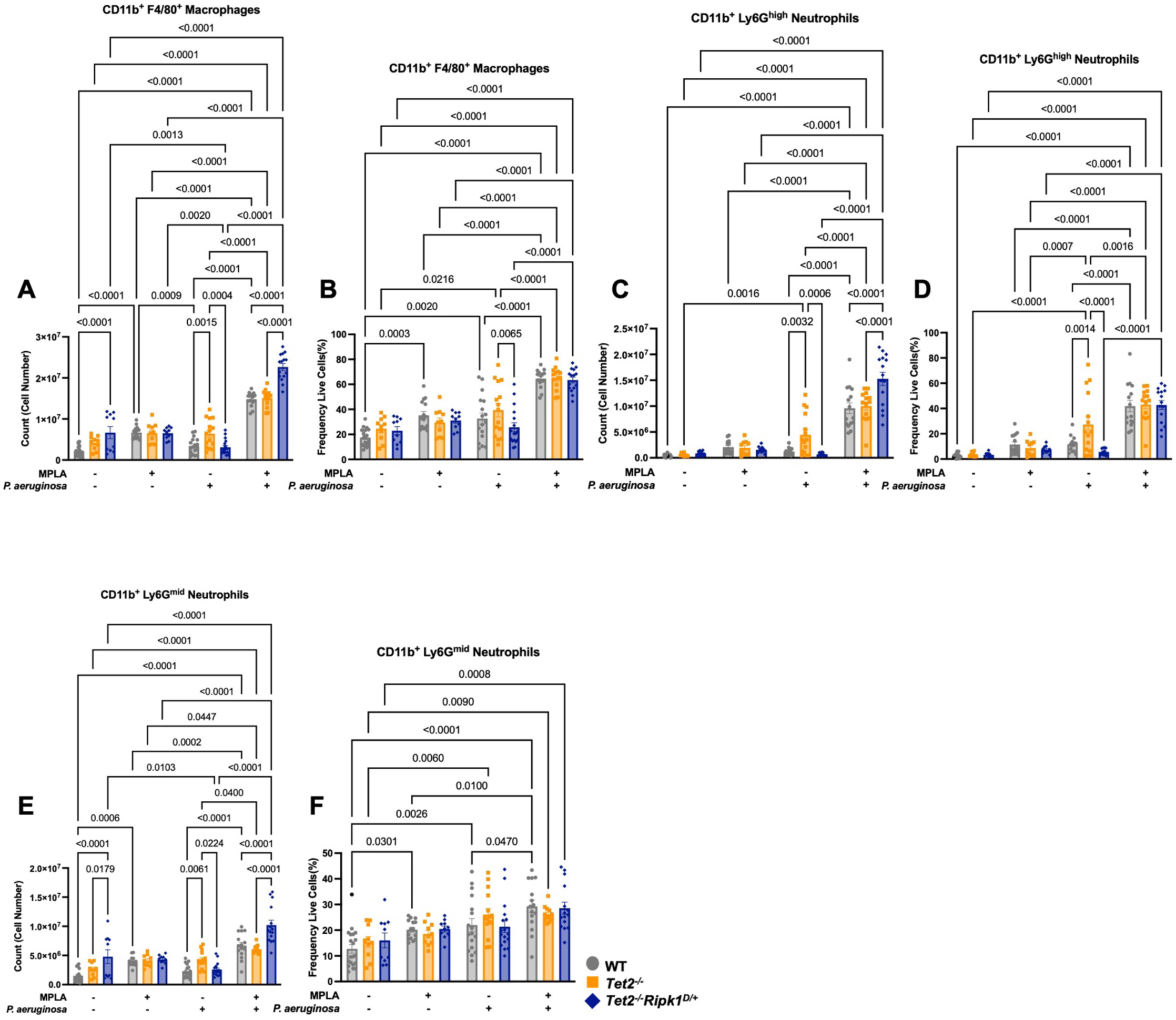
Peritoneal fluid innate immune cell populations in all treatment groups. Mice were pretreated with vehicle or MPLA for 3 days, then mock infected or infected with *P. aeruginosa* IP for 6 hours. Spectral flow cytometry was performed on peritoneal fluid, and shown are **(A)** CD11b^+^ F4/80^+^ macrophages total count and **(B)** frequency of live cells, **(C)** CD11b^+^ Ly6G^high^ neutrophils total count and **(D)** frequency of live cells, **(E)** and CD11b^+^ Ly6G^mid^ neutrophils total count and **(F)** frequency of live cells. Data are shown as mean ± SEM, 2-way ANOVA with post-hoc Tukey’s test, n = 10-20 mice per group, each point represents an individual mouse.

**Supplemental Figure 3:**
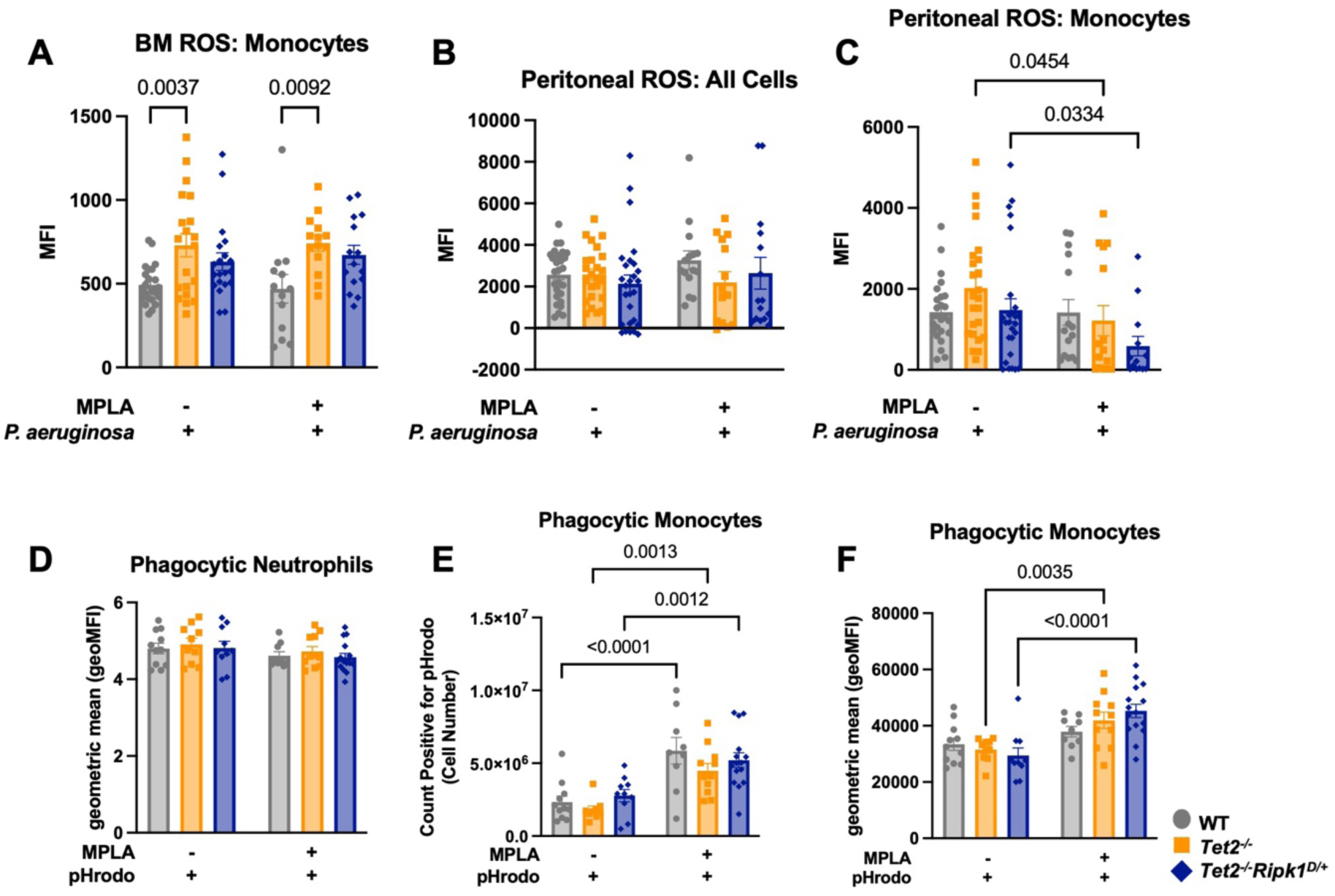
Functional capacity assessments of innate immune cells. Mice were given vehicle or MPLA 3 days prior to 6-hour infection with *P. aeruginosa*, following which whole bone marrow was collected. **(A)** Cytosolic ROS of monocytes within bone marrow as a readout of mean fluorescent intensity (MFI). Peritoneal fluid from mice was examined for ROS production, and shown are **(B)** all cell cytosolic ROS and **(C)** monocyte ROS. *In vivo* phagocytosis was determined by injecting mice with MPLA for 24 hours prior to IP injection with *E. coli* conjugated pHrodo beads, following which peritoneal fluid was harvested for assessment by flow cytometry. Shown are neutrophil (Ly6G^+^) phagocytosis as **(D)** gMFI of pHrodo, and monocyte phagocytosis (Ly6C^+^) as **(E)** cell number positive for pHrodo, and **(F)** gMFI. Data are shown as mean ± SEM, 2-way ANOVA with post-hoc Tukey’s test, n = 5-23 mice per group for bone marrow ROS, n = 10-28 mice per group for peritoneal ROS, and 13-17 mice per group for phagocytosis. Each point represents an individual mouse.

**Supplemental Figure 4:**
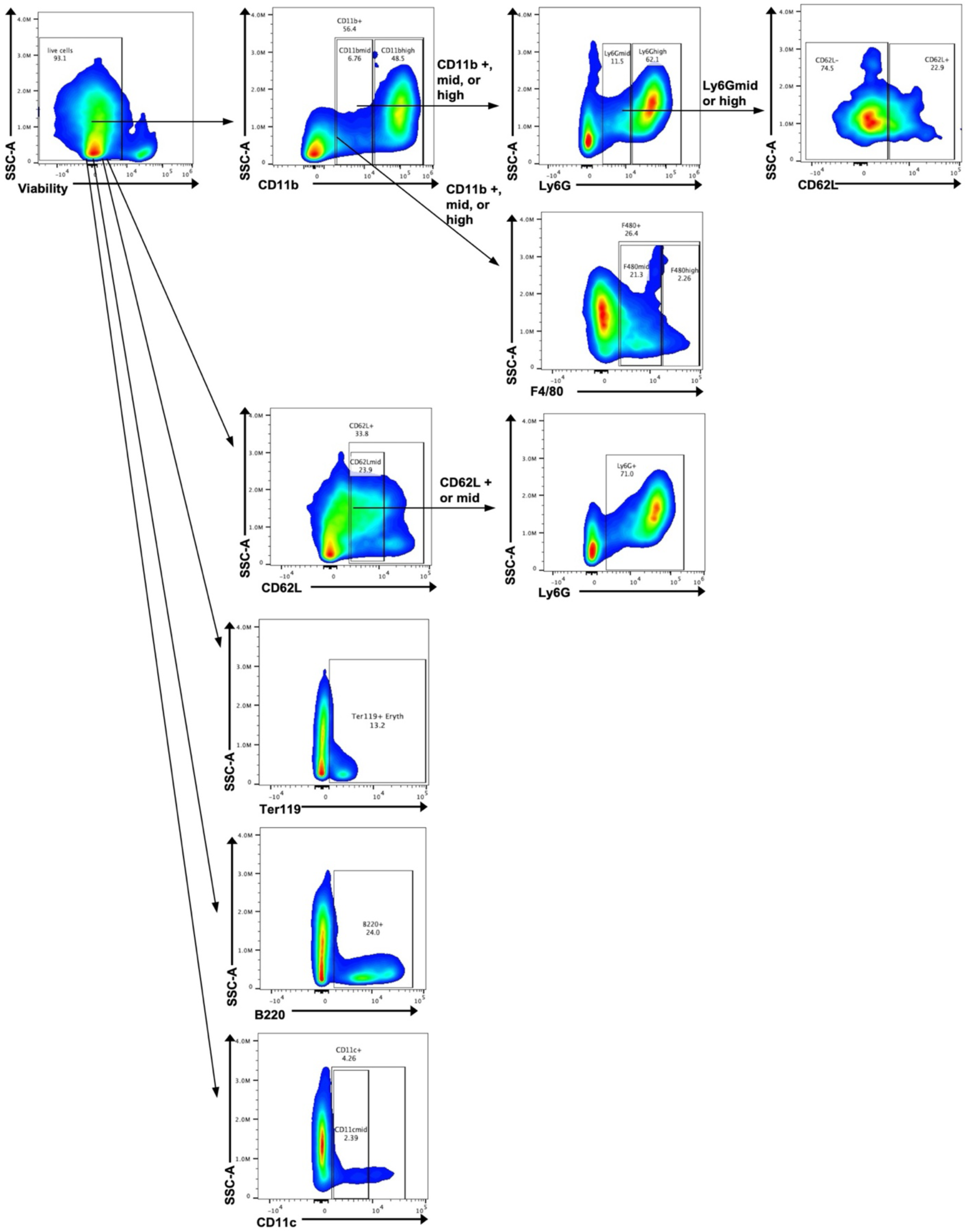
Gating strategy for mature myeloid populations via spectral flow cytometry. Live cells were gated by negativity for live/dead stain. Next, CD11b +, mid, and high populations were determined. Within these populations. Ly6G mid and high populations could be determined, as well as F4/80 +, mid, or high. Within Ly6G mid or high, CD62L – and + populations are identified. Alternatively, within the live cell population, CD62L + and mid populations can be identified, and Ly6G^+^ within either population. Additionally, within live cells, Ter119^+^ erythroid cells are identified. Lastly, live cells are gated for on B220^+^ or CD11c^+^ to identify B cell populations or dendritic cell populations, respectively.

**Supplemental Figure 5:**
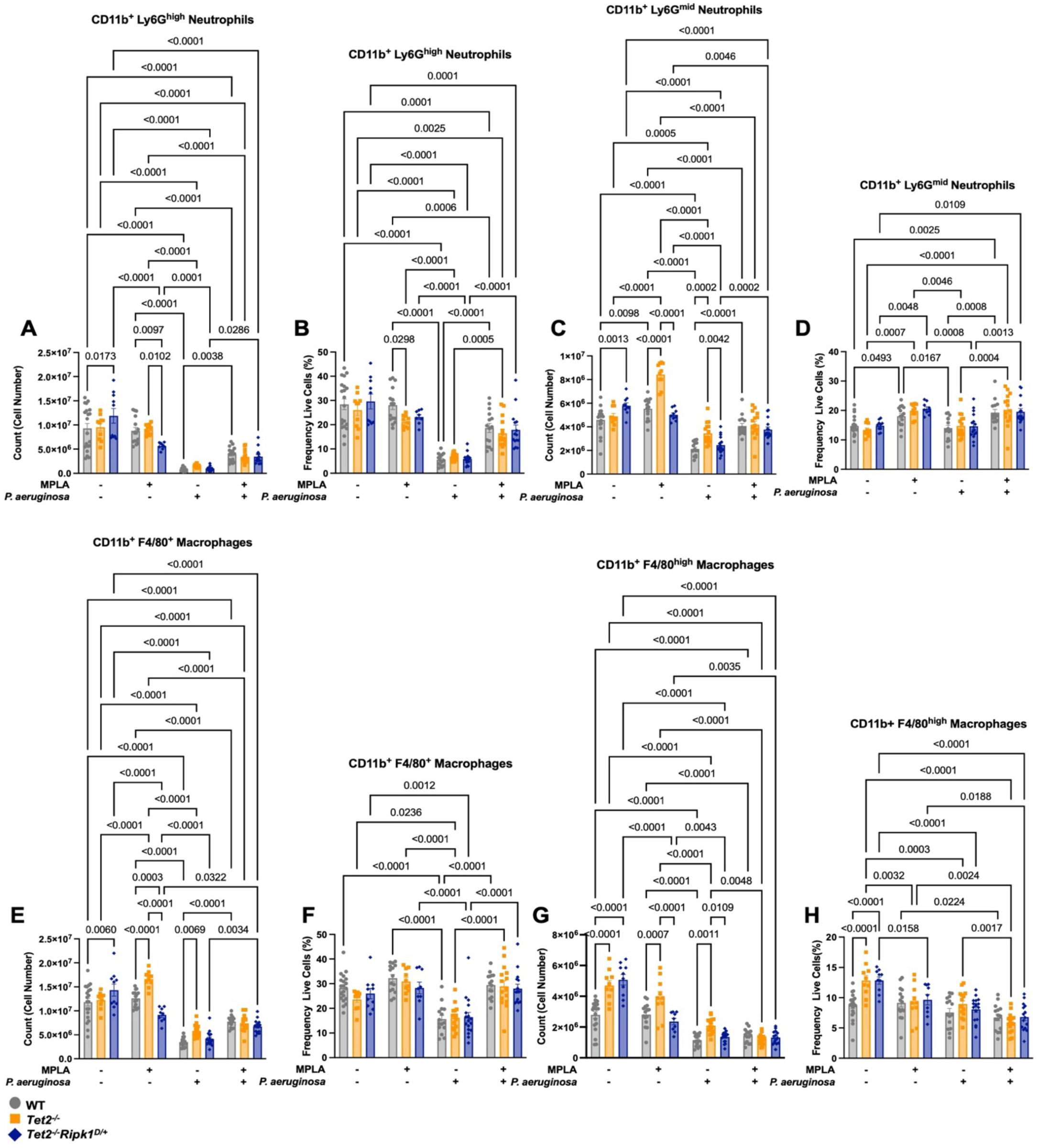
Bone marrow neutrophil and macrophage cell counts and frequencies by spectral flow cytometry. Mice were given vehicle or MPLA for 3 days and then mock infected or infected with *P. aeruginosa* for 6 hours. Then, whole bone marrow was harvested, and spectral flow cytometry was performed for neutrophil populations. Shown are **(A)** CD11b^+^ Ly6G^high^ neutrophil total counts and **(B)** frequency of live cells, representing mature neutrophils, **(C)** CD11b^+^ Ly6G^mid^ neutrophil total counts and **(D)** frequency of live cells, a less mature neutrophil population, **(E)** CD11b^+^ F4/80^+^ macrophage and **(F)** frequency of live cells, and **(G)** CD11b^+^ F4/80^high^ macrophage total cell count and **(H)** frequency of live cells. Data are shown as mean ± SEM, 2-way ANOVA with post-hoc Tukey’s test, n = 9-20 mice per group. Each mouse represents an individual mouse.

**Supplemental Figure 6:**
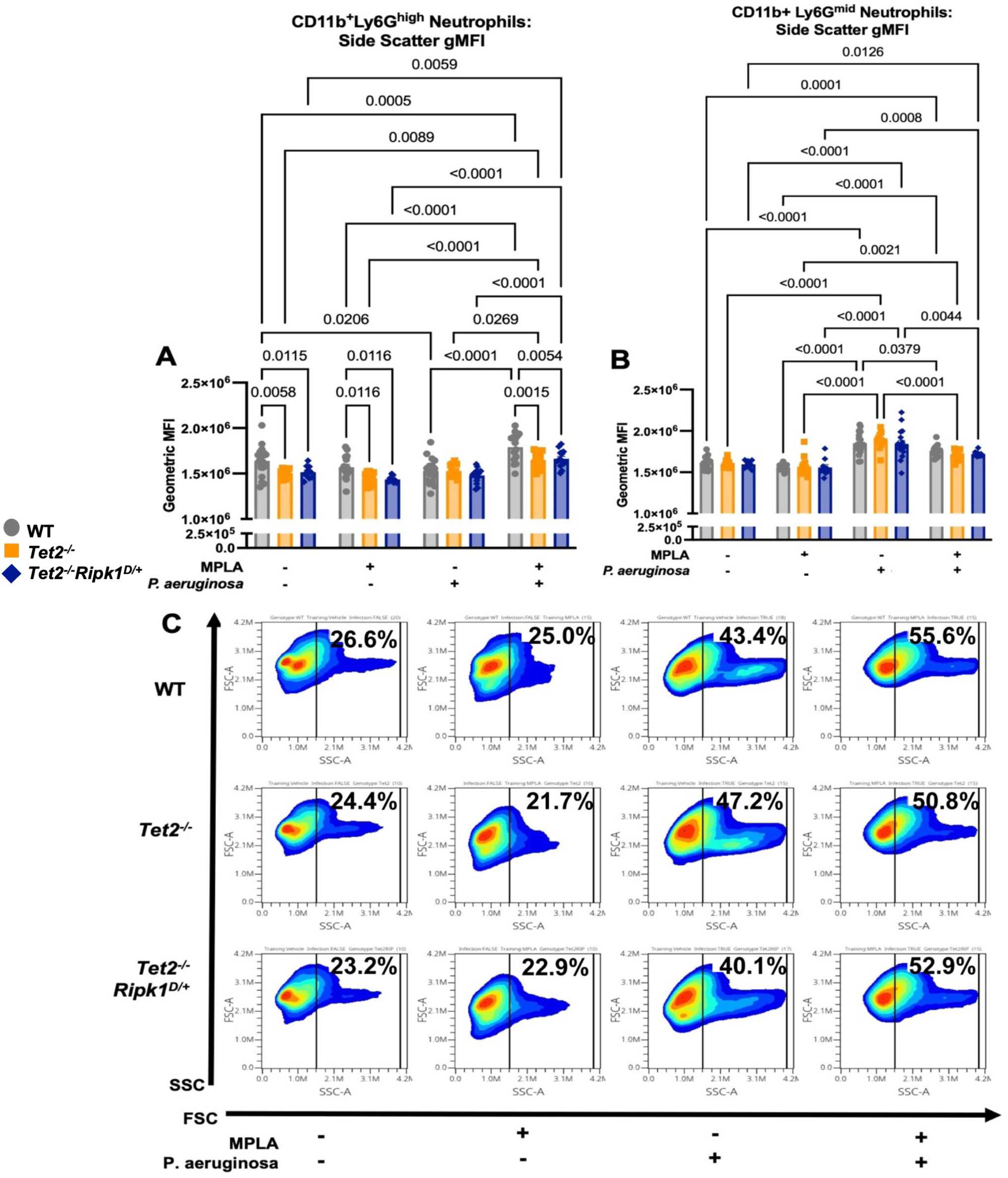
Bone marrow differentiation deficits by side scatter gMFI assessed by spectral flow cytometry. Mice were given vehicle or MPLA for 3 days and then mock infected or infected with *P. aeruginosa* for 6 hours. Whole bone marrow was harvested and assessed by spectral flow cytometry for mature and immature neutrophil populations. **(A)** Geometric mean fluorescent intensity (gMFI) for side scatter on gated CD11b^+^ Ly6G^high^ (mature neutrophils) and **(B)** CD11b^+^ Ly6G^mid^ immature neutrophils. **(C)** Combined flow plots from all treatment groups (vehicle and MPLA pretreated, and mock or *P. aeruginosa* infected). Gated immature neutrophils (CD11b^+^ Ly6G^mid^) were plotted for FSC and SSC, and frequencies of FSC are shown. Data are shown as mean ± SEM, 2-way ANOVA with post-hoc Tukey’s test, n = 9-20 mice per group. Each point represents an individual mouse.

**Supplemental Figure 7:**
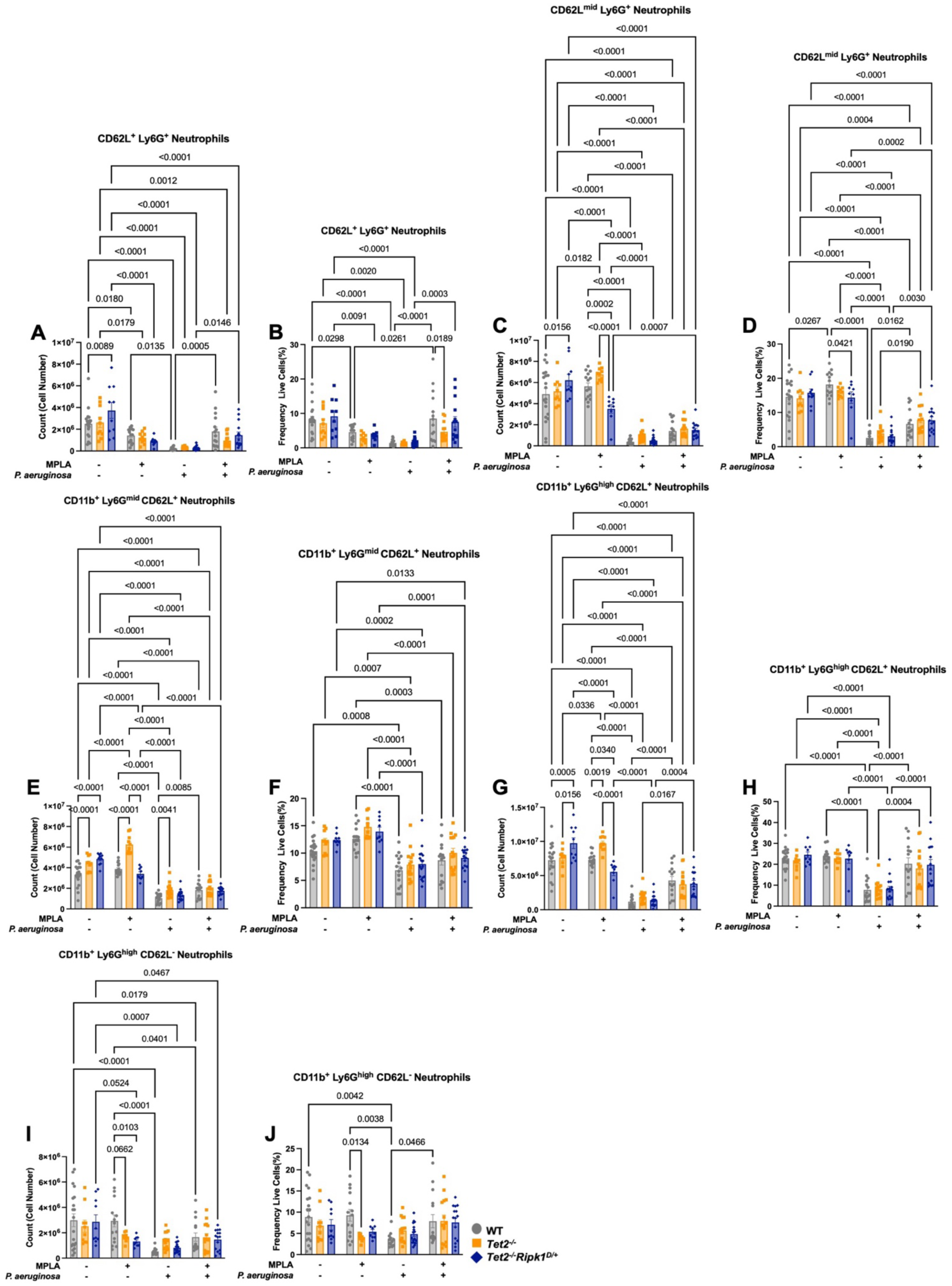
Bone marrow neutrophil population cell counts and frequencies by CD62L expression assessed by spectral flow cytometry. Mice were given vehicle or MPLA for 3 days and then mock infected or infected with *P. aeruginosa* for 6 hours. Whole bone marrow was harvested and assessed by spectral flow cytometry. Shown are **(A)** CD62L^+^ Ly6G^+^ neutrophil total cell count and **(B)** frequency of live cells, **(C)** CD62L^mid^ Ly6G^+^ neutrophil total cell count and **(D)** frequency of live cells, **(E)** CD11b^+^ Ly6G^mid^ CD62L^+^ neutrophil total cell count and **(F)** frequency of live cells, **(G)** CD11b^+^ Ly6G^high^ CD62L^+^ neutrophil total cell count and **(H)** frequency of live cells, and **(I)** CD11b^+^ Ly6G^high^ CD62L^-^ neutrophil total cell count and **(J)** frequency of live cells. Data are shown as mean ± SEM, 2-way ANOVA with post-hoc Tukey’s test, n = 9-20 mice per group. Each point represents an individual mouse.

**Supplemental Figure 8:**
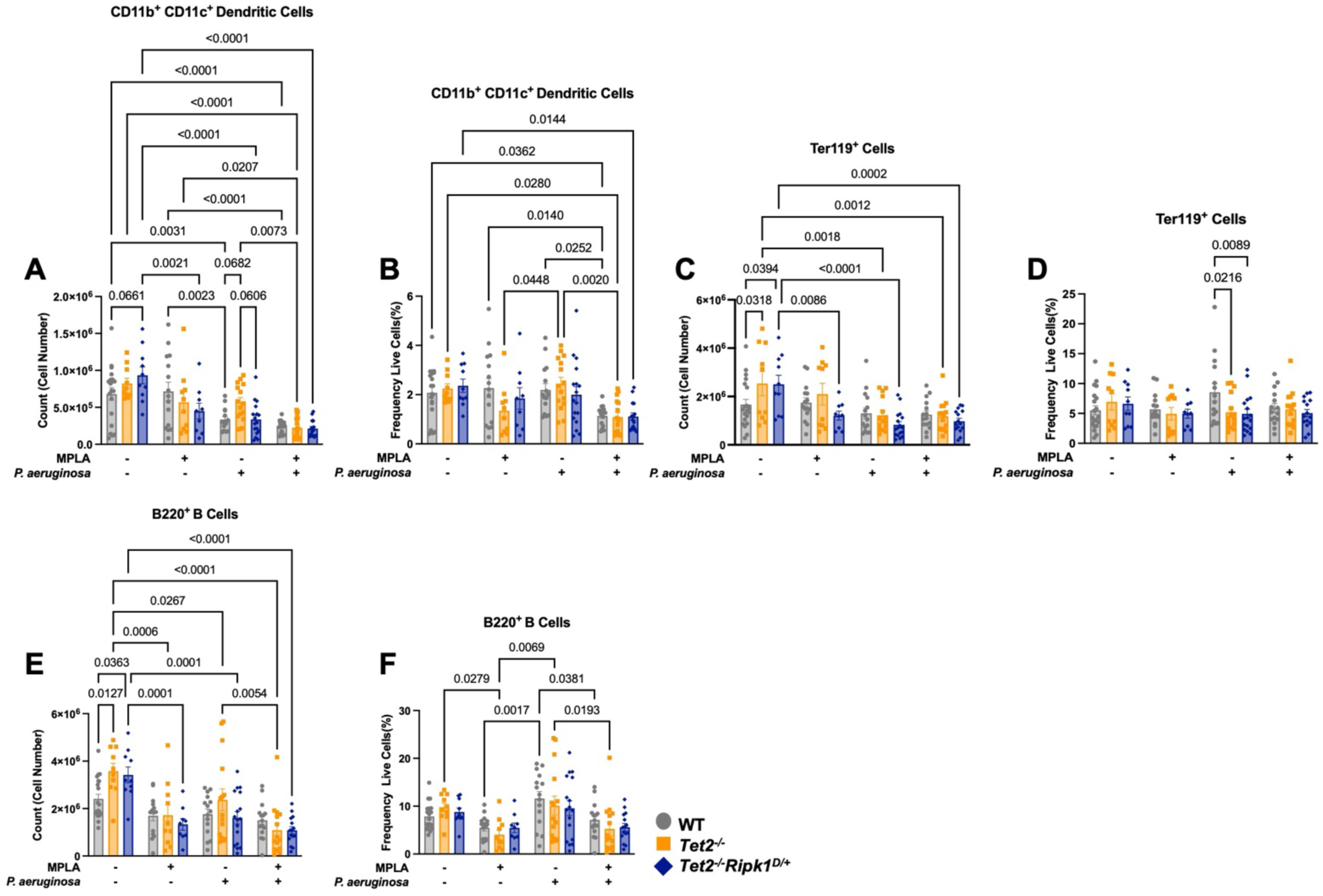
Bone marrow dendritic cell, erythroid, and B cell population counts and frequencies assessed by spectral flow cytometry. Mice were given vehicle or MPLA for 3 days and then mock infected or infected with *P. aeruginosa* for 6 hours. Whole bone marrow was harvested and assessed by spectral flow cytometry. Dendritic cell populations shown are **(A)** CD11b^+^ CD11c^+^ dendritic cell total cell count and **(B)** frequency of live cells. Erythroid populations shown are **(C)** Ter119^+^ total cell count and **(D)** frequency of live cells. Also shown are **(E)** B220^+^ B cell total cell counts and **(F)** frequency of live cells. Data are shown as mean ± SEM, 2-way ANOVA with post-hoc Tukey’s test, n = 9-20 mice per group. Each point represents an individual mouse.

**Supplemental Figure 9:**
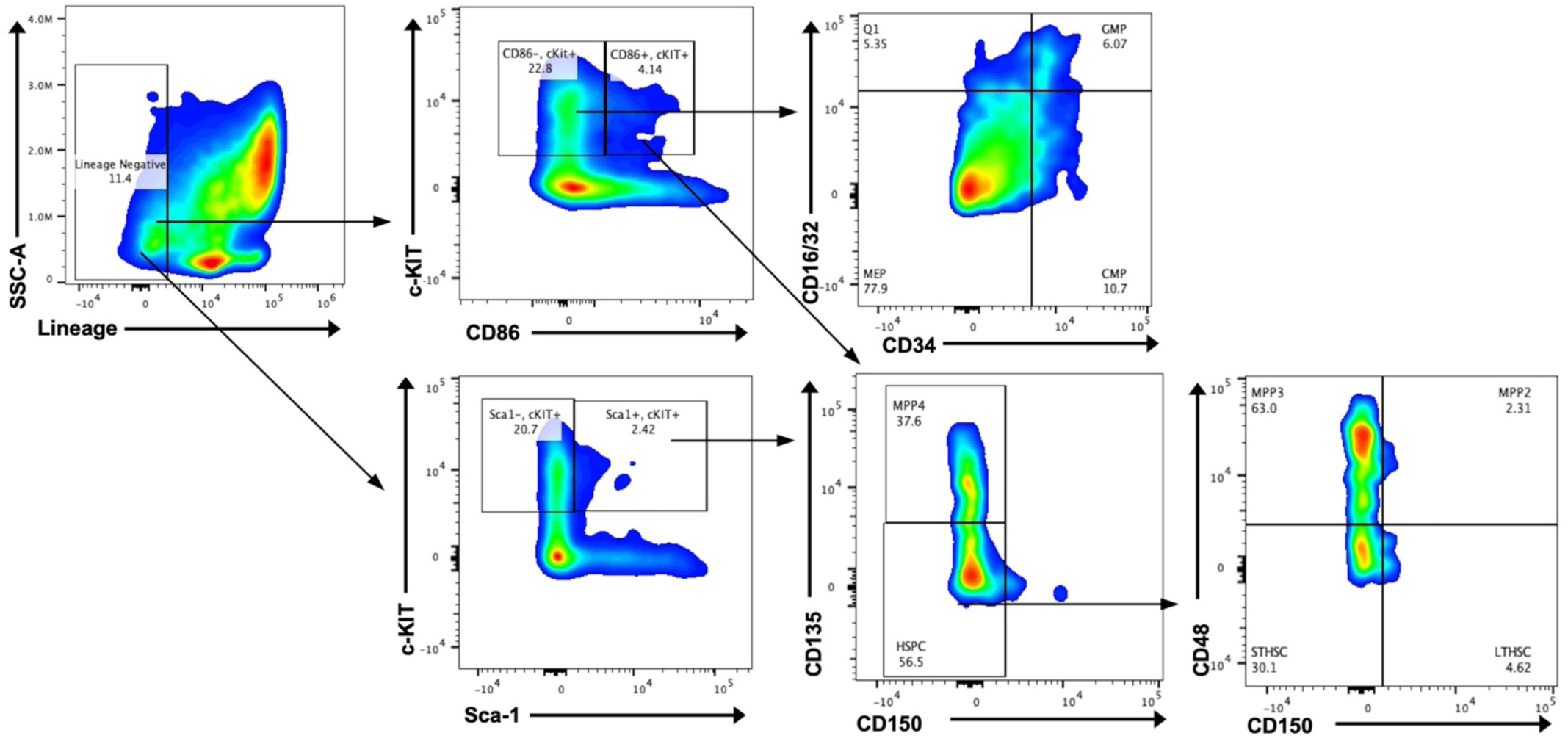
Gating strategy for bone marrow stem and progenitor populations via spectral flow cytometry. Live cells were gated for by negativity for live/dead stain. Then, lineage negative (Lin^-^) cells were gated for. Within lineage negative cells, CD86 and cKit are applied to determine CD86^+^ cKit^+^ (L86K) and CD86^-^ cKit^+^ populations. Additionally, Sca-1 and cKit can also be applied to Lin^-^ cells to determine Sca-1^+^ cKit^+^ (LSK) and Sca-1^-^ cKit^+^ populations. CD150 and CD135 is then applied to either L86K or LSK populations to determine MPP4 (CD135^+^ CD150^-^) and HSPC (CD135^-^ CD150^-^). CD150 and CD48 are then applied to HSPC to determine MPP3 (CD150^-^ CD48^-^), MPP2 (CD150^+^, CD48^+^), STHSC (CD150^-^ CD48^-^), and LTHSC (CD150^+^ CD48^-^). Additionally, CD86^-^ cKit^+^ populations are gated for by CD34 and CD16/32 to determine GMP (CD34^+^ CD16/32^+^), MEP (CD34^-^ CD16/32^-^), and CMP populations (CD34^+^ CD16/32^-^).

**Supplemental Figure 10:**
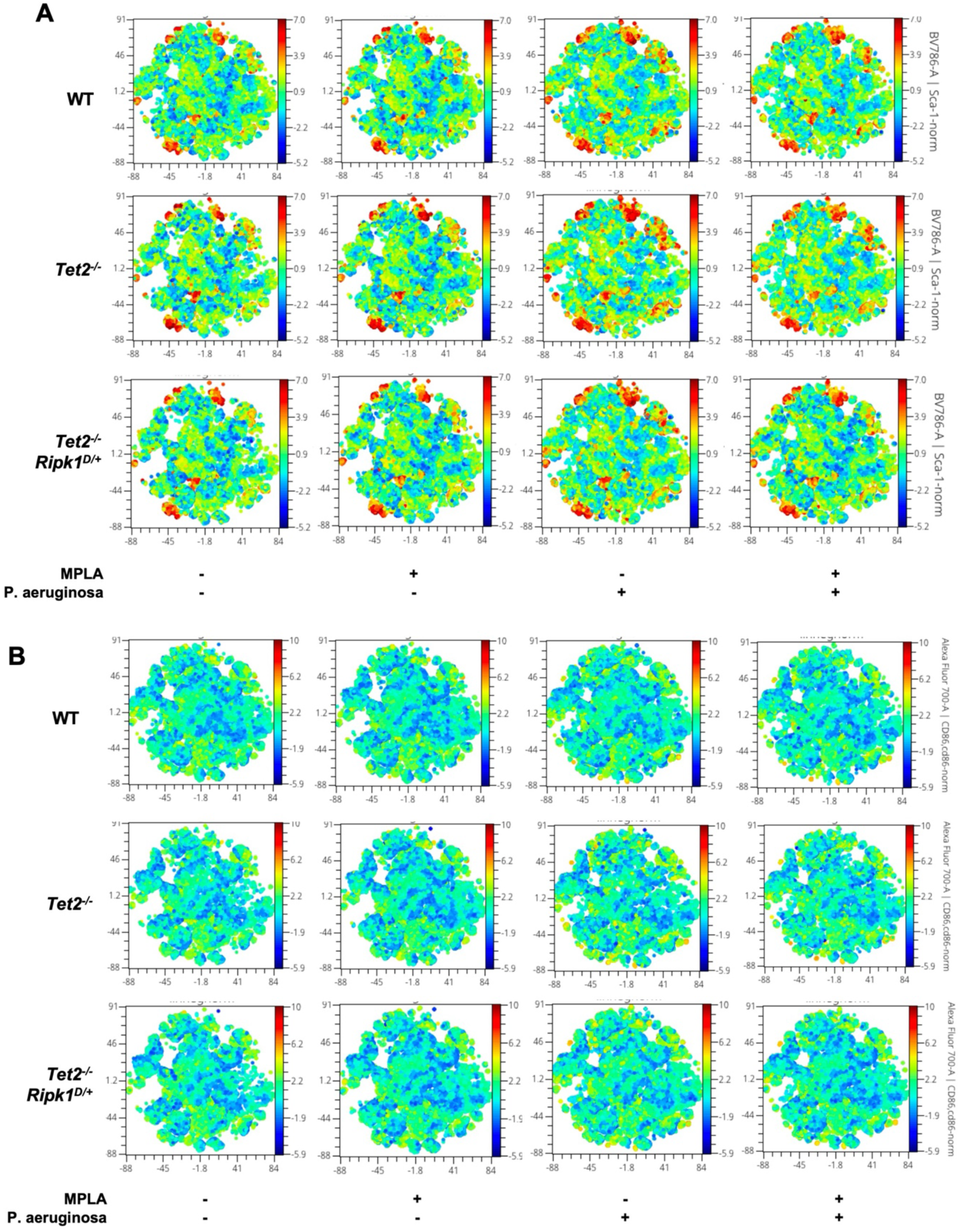
tSNE plot comparisons of Sca-1 and CD86 expression in Lin^-^ populations. Whole bone marrow from mice given vehicle or MPLA for 3 days and then mock infected or infected with *P. aeruginosa* for 6 hours. Spectral flow cytometry was performed, and Lin^-^ populations were gated from live cells. tSNE plots representing **(A)** Sca-1 expression and **(B)** CD86 expression. All biological replicates overlaid on each plot, n = 5-16 biological replicates.

**Supplemental Figure 11:**
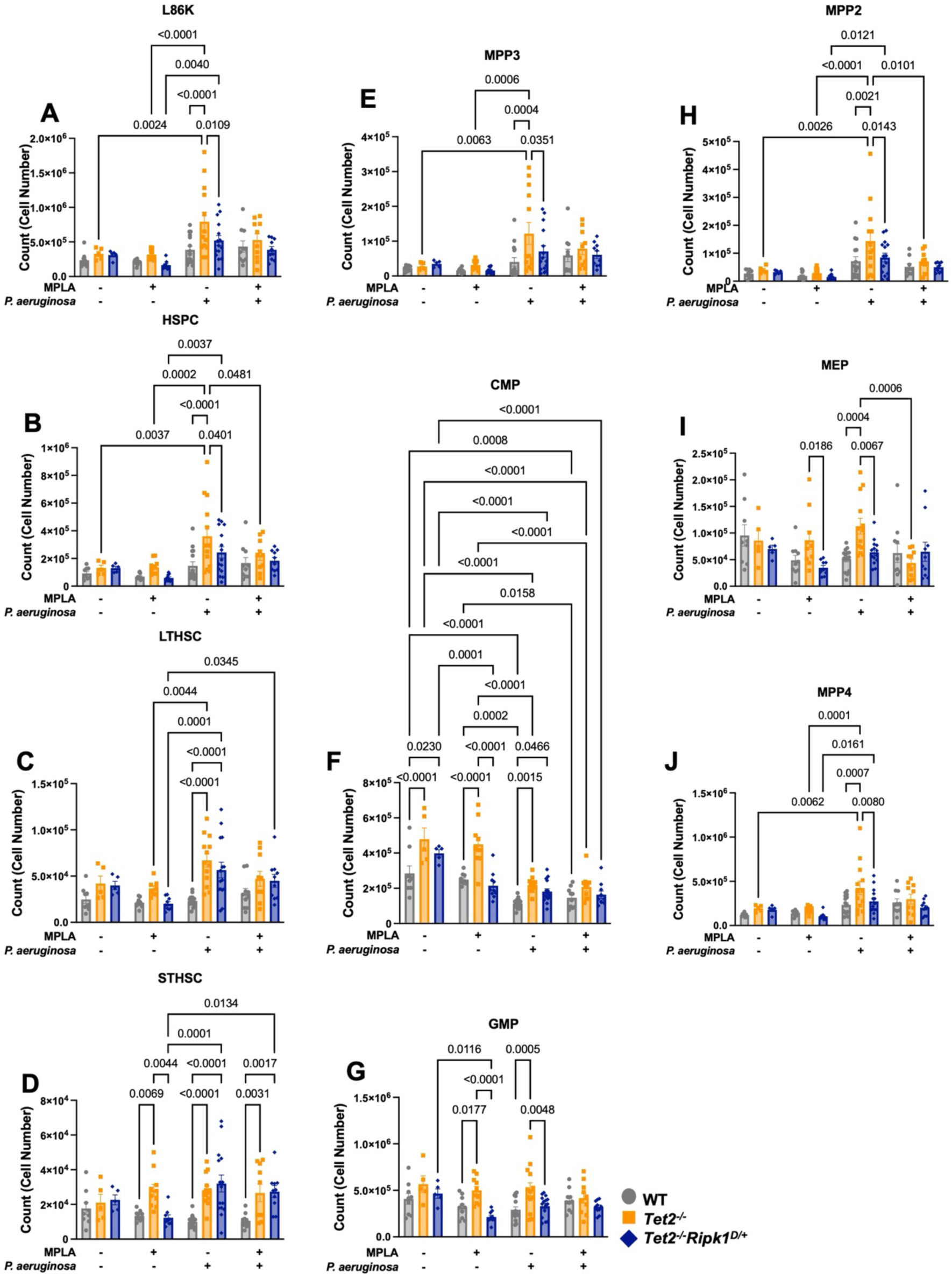
Bone marrow stem and progenitors by L86K gating strategy, total cell counts, assessed by spectral flow cytometry. Mice were given vehicle or MPLA for 3 days and then mock infected or infected with *P. aeruginosa* for 6 hours. Whole bone marrow was harvested and assessed by spectral flow cytometry. Shown are total cell counts of **(A)** L86K, **(B)** hematopoietic stem and progenitor (HSPC), **(C)** long-term hematopoietic stem cell (LTHSC), **(D)** short term HSC (STHSC), **(E)** multipotent progenitor population 3 (MPP3), **(F)** common myeloid progenitor (CMP), **(G)** granulocyte monocyte progenitor (GMP), **(H)** multipotent progenitor population 2 (MPP2), **(I)** megakaryocyte erythroid progenitor (MEP), and **(J)** multipotent progenitor population 4 (MPP4). Data are shown as mean ± SEM, 2-way ANOVA with post-hoc Tukey’s test, n = 5-16 mice per group. Each point represents an individual mouse.

**Supplemental Figure 12:**
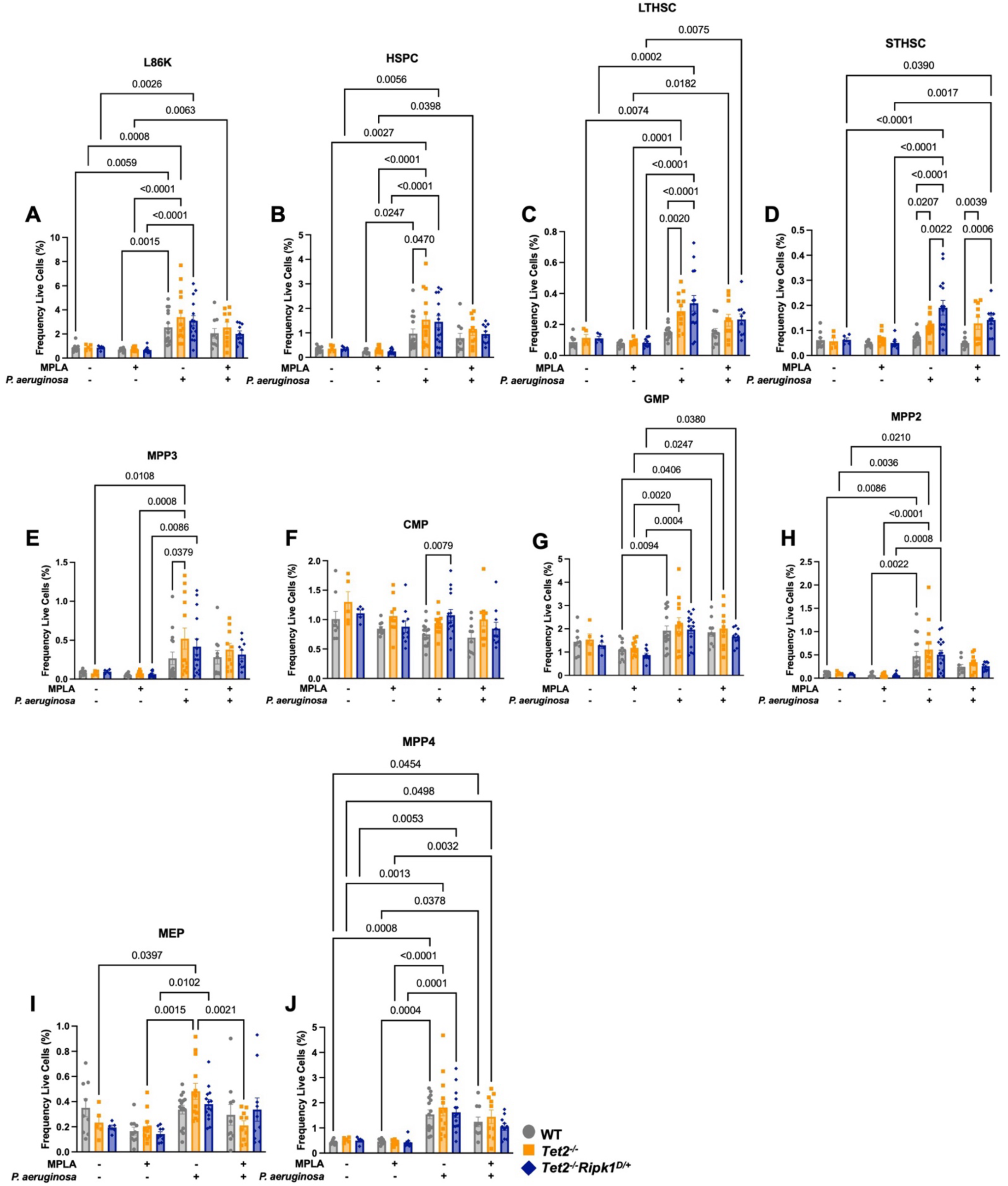
Bone marrow stem and progenitors by L86K gating strategy, frequency of live cells, assessed by spectral flow cytometry. Mice were given vehicle or MPLA for 3 days and then mock infected or infected with *P. aeruginosa* for 6 hours. Whole bone marrow was harvested and assessed by spectral flow cytometry. Shown are frequencies of live cells of **(A)** L86K, **(B)** HSPC, **(C)** LTHSC, **(D)** STHSC, **(E)** MPP3, **(F)** CMP, **(G)** GMP, **(H)** MPP2, **(I)** MEP, and **(J)** MPP4. Data are shown as mean ± SEM, 2-way ANOVA with post-hoc Tukey’s test, n = 5-16 mice per group. Each point represents an individual mouse.

**Supplemental Figure 13:**
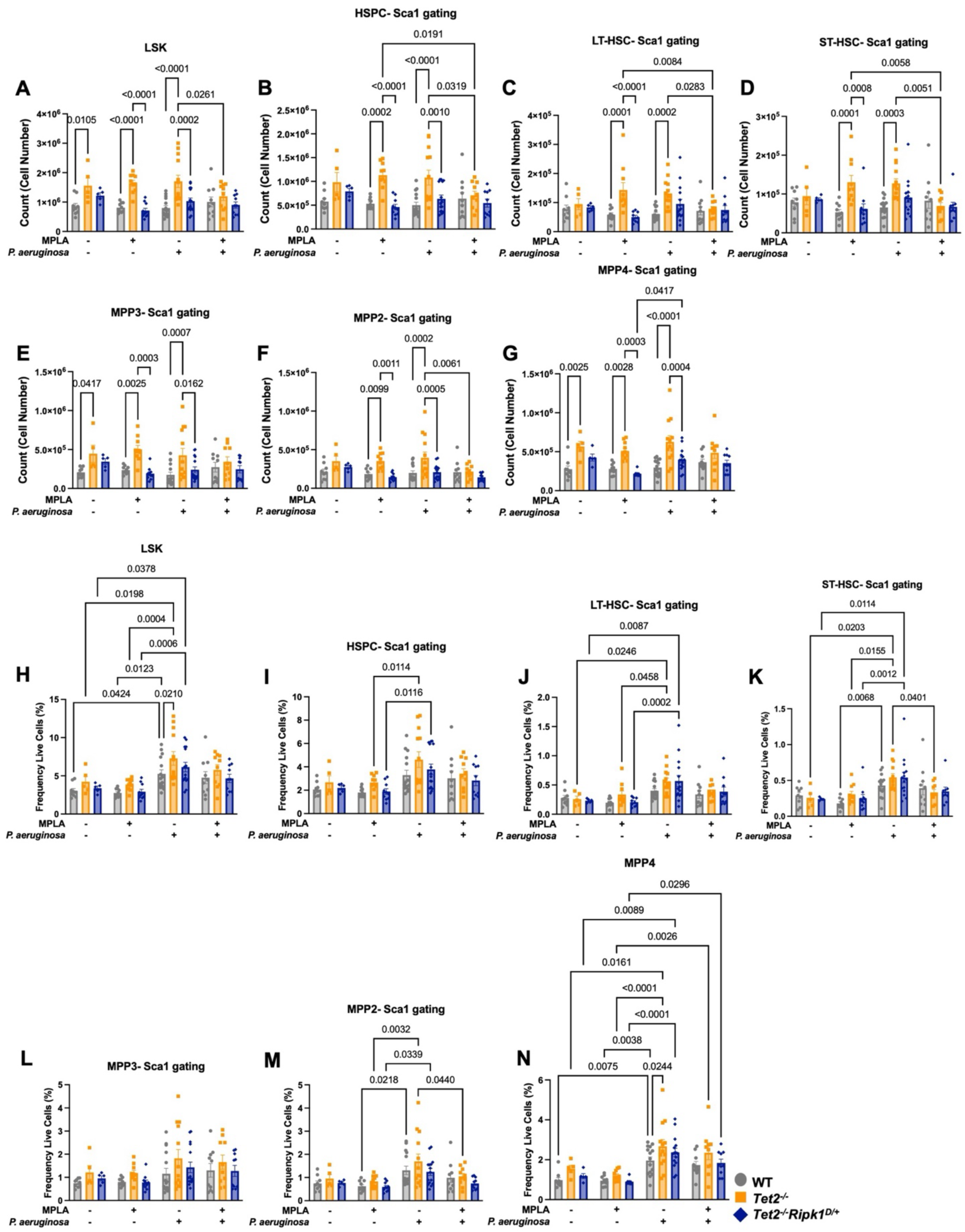
Bone marrow stem and progenitors by Sca-1 gating strategy by assessed by spectral flow cytometry. Mice were given vehicle or MPLA for 3 days and then mock infected or infected with *P. aeruginosa* for 6 hours. Whole bone marrow was harvested and assessed by spectral flow cytometry. Shown are total cell counts of (A) LSK, (B) HSPC, (C) LT-HSC, (D) ST-HSC, (E), MPP3, (F) MPP2, and (G) MPP4. Additionally shown are frequencies of total cells of (H) LSK, (I) HSPC, (J) LT-HSC, (K) ST-HSC, (L) MPP3, (M) MPP2, and (N) MPP4. Data are shown as mean ± SEM, 2-way ANOVA with post-hoc Tukey’s test, n = 5- 16 mice per group. Each point represents an individual mouse.

